# Generating dynamic structures through physics-based sampling of trRosettaX2-predicted inter-residue geometries

**DOI:** 10.1101/2025.05.28.656531

**Authors:** Chenxiao Xiang, Wenkai Wang, Zhenling Peng, Jianyi Yang

## Abstract

Deep learning methods like AlphaFold2 have achieved remarkable breakthroughs in predicting static protein structures. However, accurately modeling alternative conformations and dynamic structures remains a significant challenge. Here, we introduce trRosettaX2-Dynamics (trX2-D), a novel approach to addressing this challenge through physics-based iterative sampling of the trRosettaX2-predicted inter-residue geometric distributions. trX2-D was first pre-trained on high-resolution X-ray structures and then fine-tuned on ∼7,000 dynamic NMR structures, enhancing its inherent ability to predict alternative conformations and dynamic structures. Leveraging a transformer-based neural network, it first predicts an initial set of inter-residue geometric constraints, which are then sampled to generate dynamic structures iteratively, without requiring any prior knowledge of native structural states. Comprehensive benchmarks on three datasets, two for alternative conformations and one for dynamic structures, demonstrate that trX2-D shows promise in predicting alternative conformations and capturing structural dynamics. This work illustrates the potential of combining deep learning predictions with physics-based sampling for advancing the prediction of protein dynamic structures.

## Introduction

Protein structures are fundamental determinants of biological function^1^, and their dynamic conformational changes orchestrate many cellular processes^2^. Understanding their conformational landscape is crucial for deciphering mechanisms of biological action, developing therapeutics, and engineering novel biological molecules. While deep learning approaches, such as AlphaFold2 (AF2)^3^, RoseTTAFold^4^, trRosetta^5–8^, and ESMFold^9^, have revolutionized static protein structure prediction, accurately modeling alternative conformations of proteins remains a significant challenge.

The challenge in predicting multiple conformations arises from the scarcity of experimental data. Recently, Bryant et al. ^10^ revealed a surprisingly small number of sequence clusters (<1,000) exhibiting significant conformational diversity in the Protein Data Bank (PDB)^11^. Consequently, current data-driven methods like AF2 tend to produce conformations that closely resemble those experimentally resolved in the PDB, posing a challenge to effectively capturing the protein conformational diversity.

Several strategies have been explored to address the challenge of multi-conformation prediction. Methods leveraging contact maps of a known conformation to predict another have demonstrated the ability to generate alternative conformations^12^. However, these methods are inherently limited as they require prior knowledge of at least one native conformational state and struggle to predict more than two distinct conformations. Molecular dynamics simulations offer a physics-based approach to explore the conformational landscape of proteins^13^, but the substantial computational cost and time requirements limit their applicability to large protein systems. The AF2-based approaches, such as AF-Cluster^14^, have shown promise in generating multiple conformations by clustering and sampling diverse inputs for AF2 (e.g., multiple sequence alignment (MSA) and template^10, 14, 15^). Nevertheless, concerns regarding the true performance of these AF2-based methods have emerged due to inadequate benchmark testing and data leakage stemming from the AF2 training set ^10, 16^. To address this issue, Cfold^10^ retrained and evaluated AF2 using a meticulously constructed training-test split, and employed MSA clustering and random dropout to generate diverse conformations. Nevertheless, their results indicated that certain conformations remain elusive using MSA clustering and random dropout strategies.

Overall, AF-Cluster and other deep learning methods rely on modifying the inputs to the model in an attempt to generate multiple conformations. However, the effectiveness of these input perturbation strategies depends on highly informative inputs. For example, AF-Cluster struggles with shallow MSAs (e.g., depth < 10). Moreover, lacking direct control over the predicted structures might limit the diversity and functional relevance of the generated conformations.

Building on this observation, we introduce trRosettaX2-Dynamics (trX2-D), a novel deep learning-based approach to predict multiple conformations using an output-driven iterative sampling strategy. This method is primarily powered by a Transformer-based protein structure prediction method, trRosettaX2 (trX2), which is an improved version of trRosettaX^6^ and outperforms RoseTTAFold though using much fewer parameters and computational resources ^17^. trX2 adopts an end-to-end architecture that can simultaneously predict the 2D geometries (1 distance and 3 orientations defined in trRosetta^5^) and 3D structures. A unique property of the predicted 2D geometries lies in that they are represented as probability distributions and thus potentially encode latent information about alternative conformations. Inspired by this, trX2-D designs a heuristic module to sample diverse conformations based on the iterative sampling of the predicted 2D geometries, which allows the generation of multiple conformations without any prior information. In addition, trX2-D employs a fine-tuning strategy on the dynamic structures solved by Nuclear Magnetic Resonance (NMR) experiments to improve the conformational diversity information in the predicted geometries.

We evaluated trX2-D on three datasets non-redundant to its training set, including two established benchmarks for dual-conformation proteins and a dataset of dynamic proteins. Benchmark tests show that trX2-D significantly improves upon the performance of the base trX2 model and shows promise in predicting alternative conformations on dual-conformation benchmarks. Furthermore, our tests on the dynamic protein dataset indicate that trX2-D can generate more diverse conformation ensembles compared to other methods like AF-Cluster. In summary, trX2-D represents a novel and promising approach for predicting protein alternative conformations, marking a solid step towards a more comprehensive understanding of protein structural dynamics.

## Results

### Overview of the Method

trRosettaX2 (trX2) is a lightweight protein structure prediction algorithm designed to achieve competitive performance using limited computational resources, which has been briefly introduced before^17^. As shown in Fig. 1a, trRosettaX2 employs a Transformer-based neural network, trFormer, to predict 2D geometries (distance and orientations) from multiple sequence alignment (MSA). The 3D structure is then folded through either structure module (i.e., end-to-end prediction) or energy minimization (i.e., two-step prediction). Although the accuracy of trX2 still slightly lags behind that of AlphaFold2 (AF2), its unique advantages, such as rapid MSA selection and the generation of decoys complementary to the AF2 predictions, helped our group win the championship in CASP15 ^17, 18^ and CASP16 experiments (https://predictioncenter.org/casp16/zscores_final.cgi). For the detailed methodology description and performance analysis of trX2, please refer to Text S1.

**Fig. 1.**
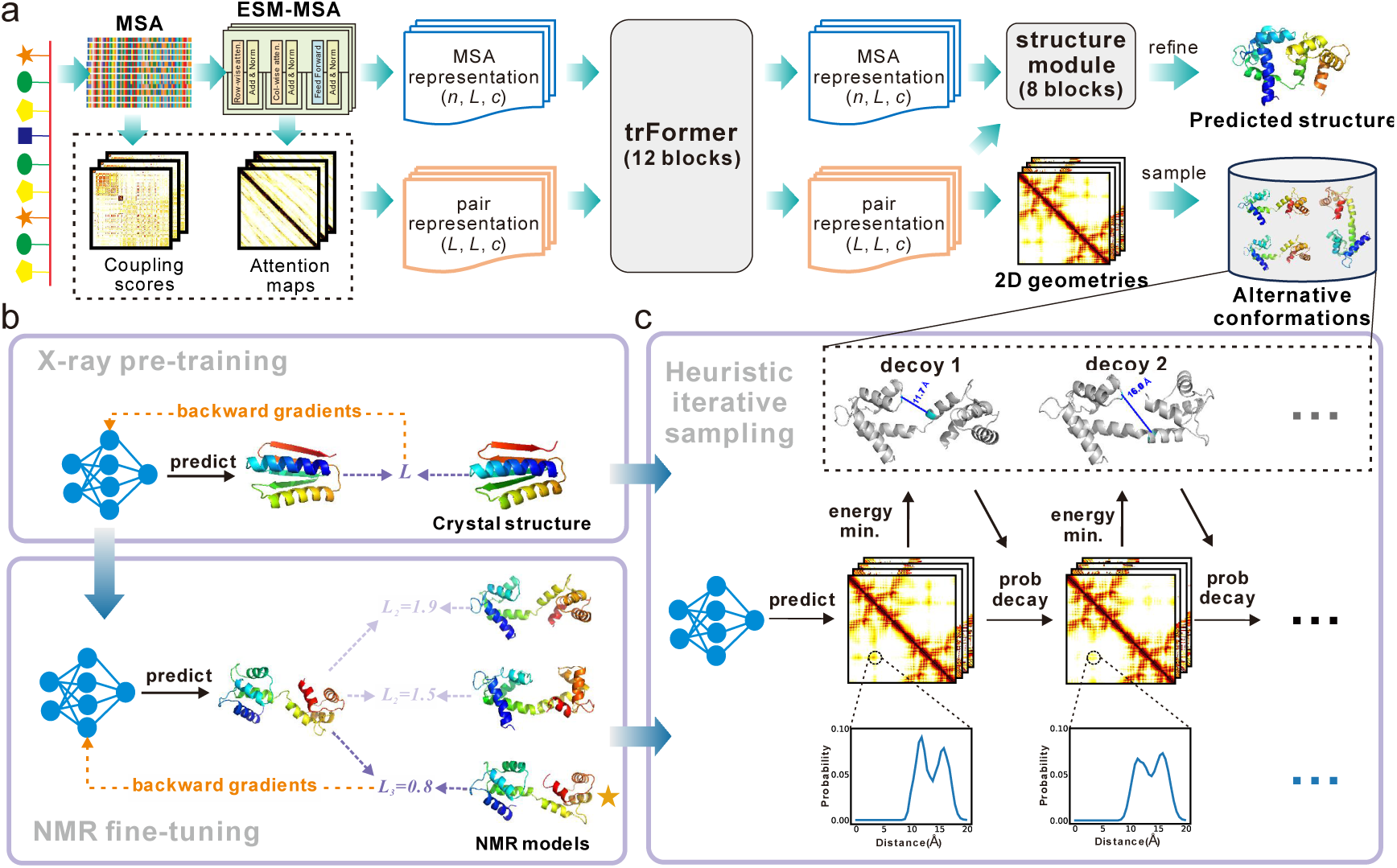
Architectures of trRosettaX2 and trRosettaX2-Dynamics. **(a)** overview of trRosettaX2 and trRosettaX2-Dynamics. The sole input is the amino acid sequence of a target protein. A multiple sequence alignment (MSA) is generated and converted into two representations, MSA representation and pair representation, which are updated through a transformer-based module (trFormer). The updated representations are fed into the structure module to predict the static 3D structure by trRosettaX2. Meanwhile, 2D geometries derived from the pair representation are used to sample alternative conformations by trRosettaX2-Dynamics. **(b)** trRosettaX2-Dynamics is first pre-trained with X-ray structures and then fine-tuned with NMR structures. **(c)** the iterative sampling of alternative conformations using predicted 2D geometries.

Building upon trRosettaX2, we developed trRosettaX2-Dynamics (trX2-D) to improve protein conformation generation. This advancement incorporates two principal modifications: 1) fine-tuning trX2 with NMR ensembles (trX2 (NMR); see Fig. 1b and Methods for details) to enhance the representation of dynamic signals within the model outputs; and 2) designing an iterative process for sampling diverse 2D geometries to generate multiple distinct conformations (see Fig. 1c, Supplementary Fig. S2, and Methods for details). The complete trX2-D workflow leverages both the original trX2 and the NMR fine-tuned trX2 (NMR) in parallel to produce two sets of initial 2D geometry predictions. These predictions subsequently serve as inputs to the iterative process, yielding a diverse ensemble of protein conformations.

### Performance of trX2-D in distinguishing apo and holo states across 38 proteins

We evaluated the performance of trX2-D using an elaborately collected dataset of 91 proteins, which were experimentally solved in apo-holo states^12, 19^. To focus our analysis on substantial conformational changes, subsequent detailed analyses centered on a subset of 38 proteins exhibiting large conformational changes (see Methods for details) ^10, 12^. The information of these conformation pairs is listed in Table S2. The results for the remaining samples are detailed in Supplementary Data, which lead to similar conclusions.

We first compare trX2-D with the default trX2 model to examine the extent of improvement in capturing alternative conformations. In this work, we use RMSD as the primary evaluation metric, which better reflects local structural variation than the TM-score. A supplementary TM-score comparison, consistent with the RMSD findings, is provided in Fig. S3. Fig. 2a illustrates the overall RMSD distribution of trX2-D and trX2 predictions for both apo and holo states. The default trX2 produces higher average RMSDs (> 5 Å) for both apo and holo state predictions. As shown in Fig. S4, trX2 predictions exhibit similar RMSD values relative to both the apo and holo states, suggesting they may represent intermediate states that deviate from the apo and holo conformations. In contrast, by sampling diverse 2D geometries, trX2-D demonstrates the capability to transition from these trX2-predicted intermediate states towards either the apo or holo state, consequently yielding enhanced predictive stability. As a result, trX2-D achieved significantly lower RMSD values (a 20%∼30% reduction) compared to trX2 for both states, highlighting its effectiveness in improving the alternative conformation generation.

**Fig. 2.**
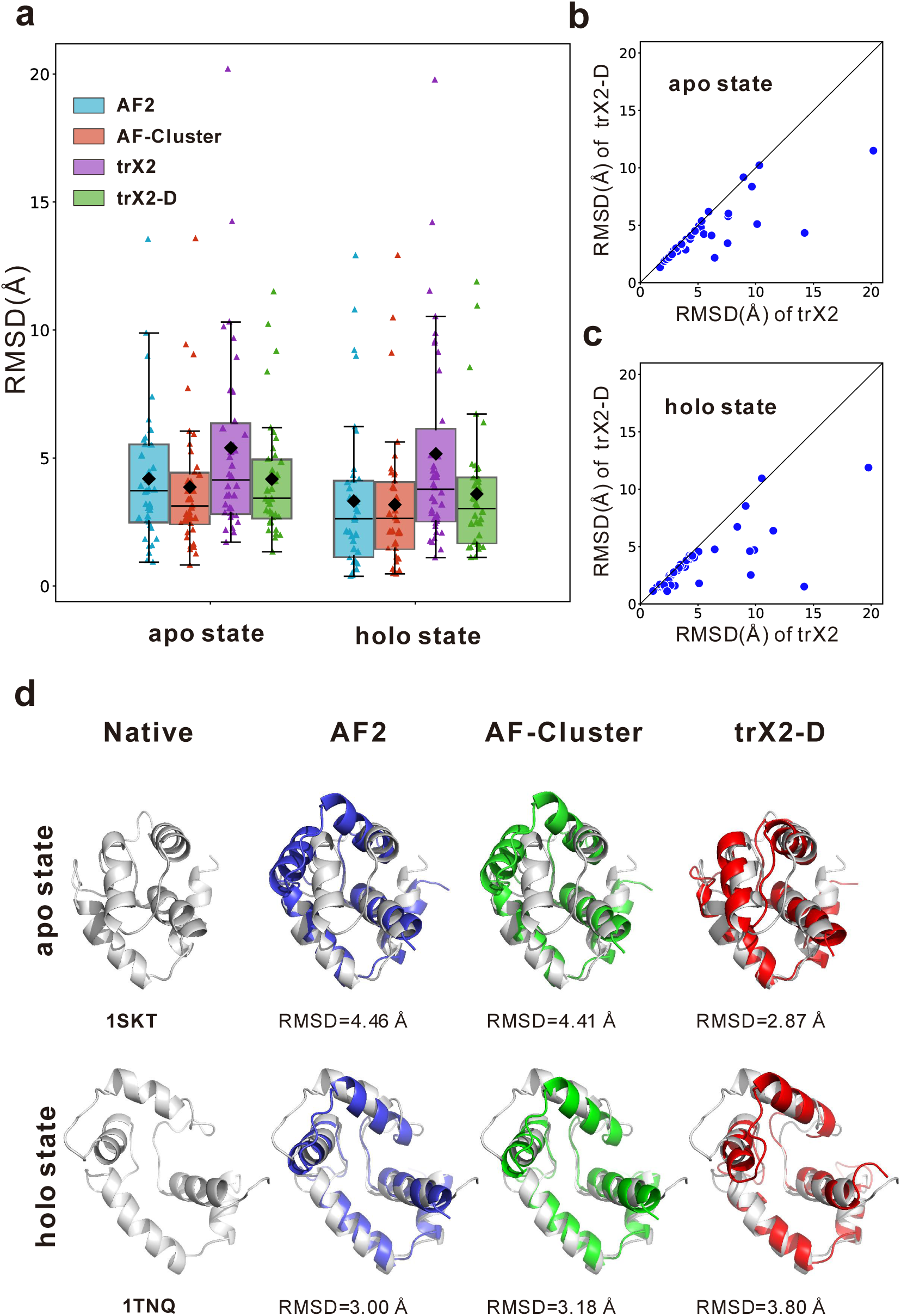
Comparative assessment of trX2-D on apo-holo state proteins. **(a)** RMSD distributions of predictions from AF2, AF-Cluster, trX2, and trX2-D against native structures, separated by apo and holo states. Individual predictions are shown as triangles. Mean RMSDs are indicated by black diamonds. **(b, c)** head-to-head RMSD comparison of trX2-D versus trX2 for the apo (b) and holo (c) states. **(d)** performance on a case with a large conformational change. For this case, the native apo (PDB ID: 1SKT) and holo (PDB ID: 1TNQ) conformations exhibit significant differences (TM-score_apo-holo_ = 0.54, RMSD_apo-holo_=7.2 Å). AF2 and AF-Cluster predict highly similar structures for the apo and holo states (TM-score_apo-holo_ ∼ 0.95, RMSD_apo-holo_ ∼ 2.0 Å). Conversely, trX2-D provides a more accurate apo prediction, and captures greater divergence between its predicted conformations for the two states (TM-score_apo-holo_ = 0.71, RMSD_apo-holo_ = 2.9 Å).

The benefits of trX2-D are further underscored by a direct head-to-head RMSD comparison with trX2 for both apo and holo states (Figs. 2b and 2c, respectively). A clear majority of data points (35/38 for the apo state; 34/38 for the holo state) fall below the diagonal line, which means that trX2-D achieves lower RMSD for ∼ 90% of samples in both states. This trend is particularly pronounced for samples where the original trX2 performed poorly (RMSD > 5 Å). Beyond accuracy, trX2-D also boosts conformational diversity. While the two conformations from trX2 were often very similar (RMSD < 2 Å for 26/38 proteins), trX2-D significantly increased their distinctiveness, with this high similarity observed in only about one-third of cases (14/38).

To gain further insight into the factors underlying this enhanced performance, we analyzed the influence of conformational diversity on the prediction accuracy of the trX2 model. Our analysis revealed that the RMSD of trX2 predictions positively correlates with the divergence between native states (denoted as RMSD_apo-holo_), with a Pearson correlation coefficient (PCC) of 0.28 (blue line in Fig. S5). This finding indicates that heterogeneity between native states tends to pose challenges to trX2’s accurate prediction. This trend is more pronounced for 16 samples exhibiting conformational differences of RMSD_apo-holo_ > 5 Å, where the PCC increases to 0.84 (orange line in Fig. S5). This highlights trX2’s difficulty in accurately modeling cases with significant structural variability. Despite this challenge faced by trX2, trX2-D achieves consistent improvements across all levels of conformational divergence (Fig. S6). This result further confirms the robust improvement made by trX2-D.

### Impact of NMR-based fine-tuning and iterative sampling

trX2-D leverages the architecture of the trX2 network. To enhance its ability to generate diverse conformations, we employed fine-tuning on a dataset of dynamic structures derived from NMR experiments^20^. Moreover, trX2-D introduces a heuristic iterative sampling process to generate diverse conformations from the predicted geometric restraints. To systematically evaluate their contributions, we conducted a series of ablation experiments, as summarized in Fig. 3 and Table S3. As a point of reference, the original trX2 produced models with average RMSD values of 5.40 Å for the apo state and 5.16 Å for the holo state, setting a baseline for assessing performance improvements.

**Fig. 3.**
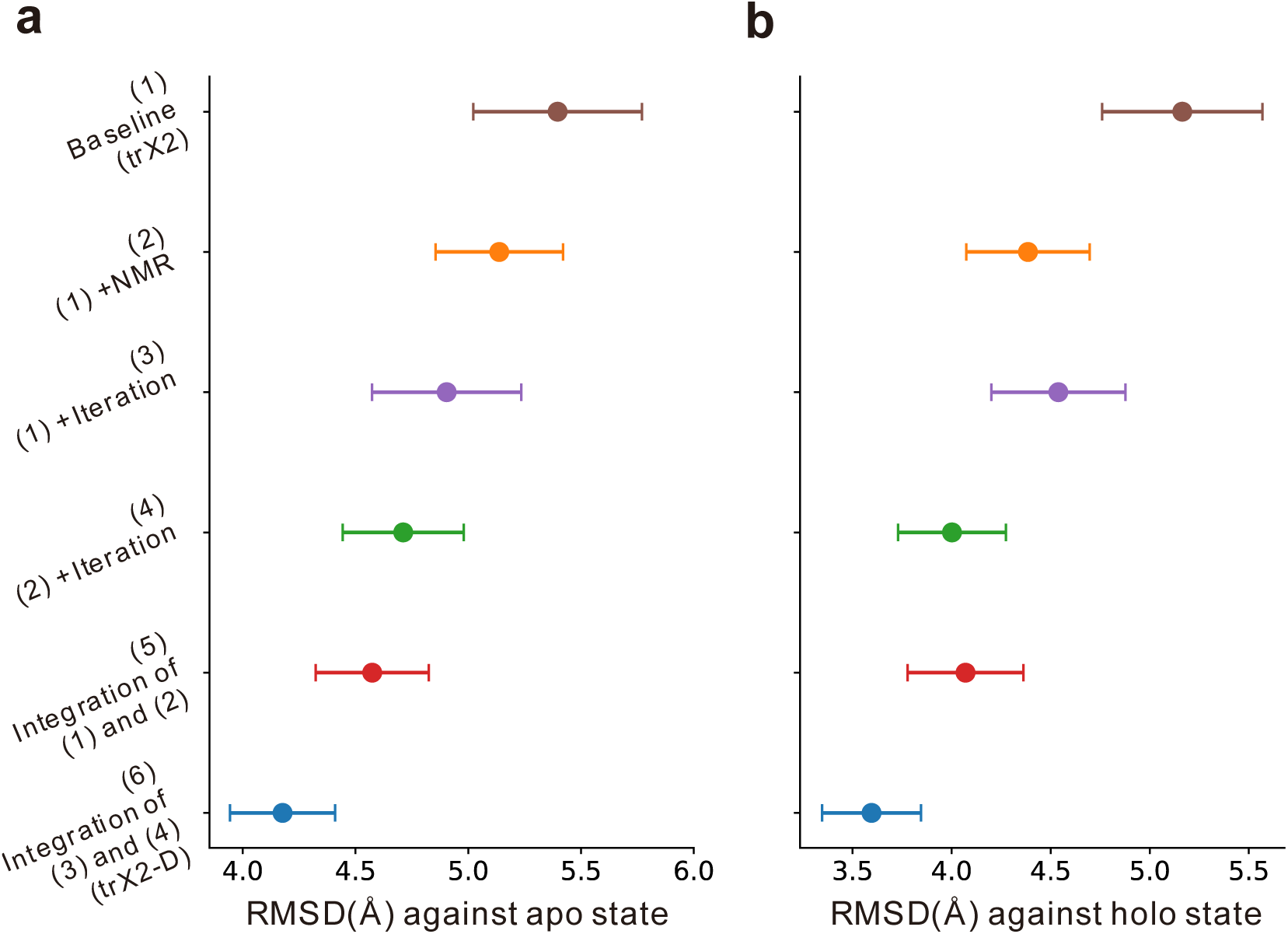
Ablation study results of trX2-D on 38 apo-holo proteins. **(a, b)** average RMSD of predicted conformations for apo (a) and holo (b) states. Error bars represent one-tenth of the standard deviation across samples. The ablation study reveals the incremental performance improvements achieved by incorporating NMR-based fine-tuning and iterative sampling, explaining the superior performance of trX2-D.

Building on the baseline model, we first evaluated the impact of NMR-based fine-tuning, which produced trX2 (NMR) (model (2) in Fig. 3) with average RMSDs of 5.13 Å (apo) and 4.38 Å (holo), both lower than trX2. This improvement suggests that fine-tuning with NMR data provides structural diversity beyond that captured by the original trX2, thereby enhancing its ability to predict multiple conformations. Interestingly, we observed that both the fine-tuned variant and the original trX2 networks demonstrated distinct advantages on certain samples. As illustrated in Fig. 4a and Fig. 4b, trX2 (NMR) outperformed trX2 for nearly half of the targets (blue points; 15/38 for the apo state, 18/38 for the holo state), likely benefiting from the additional dynamic information inherent in NMR ensembles. Conversely, trX2 (NMR) performed worse than trX2 for other targets (yellow points). This could be attributed to the noise introduced by the lower resolution and uncertainty associated with NMR structures, which might negatively impact the training data quality. These results highlighted a strong complementarity between trX2 and trX2 (NMR), indicating the potential of integrating both models to achieve more accurate multi-conformation predictions.

**Fig. 4.**
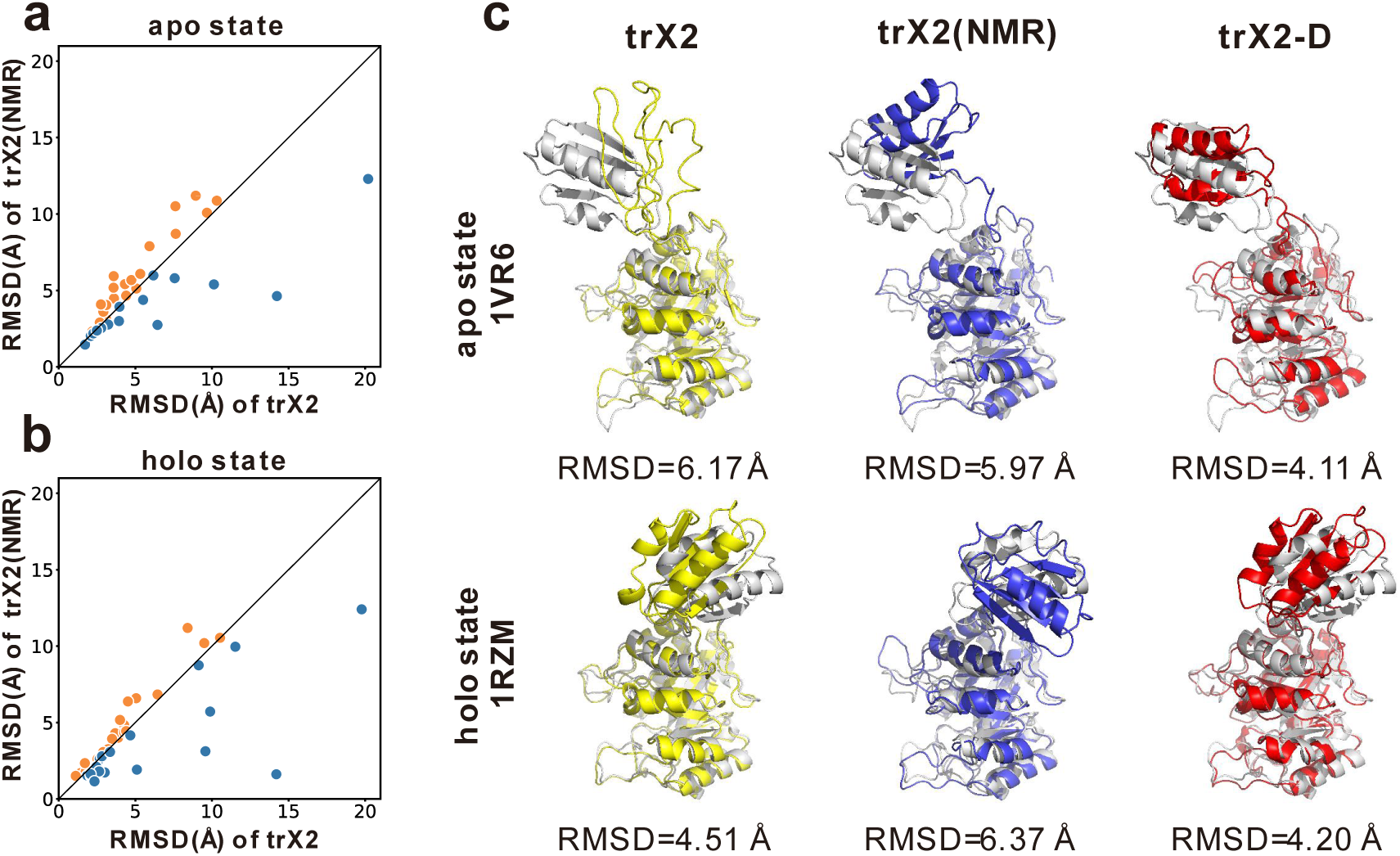
Impact of NMR-based fine-tuning. **(a, b)** head-to-head RMSD comparison between trX2 (NMR) and trX2 for the apo state (a) and the holo state (b). Orange points denote samples where trX2 performs better, while blue points indicate where trX2 (NMR) is superior. **(c)** predictions of a protein with significant inter-domain rotation (apo PDB ID: 1VR6, holo PDB ID: 1RZM). Compared to trX2, trX2 (NMR) improves apo state prediction but compromises holo state accuracy. In contrast, trX2-D improves predictions for both apo and holo states, and effectively captures the domain rotation.

Subsequently, we assessed the impact of the heuristic iterative sampling process. Employing this process to trX2 (i.e., model (3) in Fig. 3) reduced the average RMSD from 5.40 Å to 4.90 Å for the apo state, and from 5.16 Å to 4.53 Å for the holo state. Furthermore, applying the heuristic iterative process to trX2 (NMR) (i.e., model (4)) also reduced the RMSD from 5.13 Å to 4.71 Å (apo) and from 4.38 Å to 4.00 Å (holo). These results demonstrate the effectiveness of the sampling process for both trX2 and its NMR-based variant, highlighting its broad applicability. These consistent performance improvements underscore the efficacy of this sampling process in exploring conformational landscapes.

Considering the complementary modeling potential of trX2 and trX2 (NMR), we analyzed the benefits of integrating predictions from both. The integration of these two models (i.e., model (5) in Fig. 3) achieved average RMSD values of 4.57 Å (apo) and 4.07 Å (holo), surpassing single-model predictions. Building on these promising results, trX2-D (model (6) in Fig. 3) further combined the predictions from both model (3) and model (4) (i.e., the models equipped with the heuristic iterative sampling). This comprehensive integration yielded the best overall performance, with average RMSD values of 4.18 Å (apo) and 3.60 Å (holo), representing improvements of 22.6% and 30.2% compared to trX2, respectively. This strategy effectively harnesses the strengths of both models while mitigating their individual limitations. For a more specific illustration of the distinct impacts of NMR-based fine-tuning and the heuristic iterative sampling process, we analyzed a challenging test case characterized by a significant inter-domain conformation change (apo PDB ID: 1VR6, holo PDB ID: 1RZM). This protein is *3-deoxy-D-arabino-heptulosonate-7-phosphate synthase* (DAHPS) ^21^, a key enzyme for aromatic amino acid biosynthesis. Its activity is regulated by an allosteric transition between inactive and active forms upon ligand binding (Cd²⁺, PEP, and E4P), reflected as a substantial inter-domain motion involving a ∼160 ° rotation (RMSD_apo-holo_ = 10.10 Å). This pronounced conformational difference makes DAHPS a particularly challenging and informative test case for methods aiming to capture conformational diversity.

As shown in Fig. 4c, while trX2 generates a structure approximating the holo state (RMSD = 4.51 Å), it fails to accurately model the secondary structure of the variable domain in the apo state, yielding a higher RMSD of 6.17 Å. In contrast, trX2 (NMR) correctly models the secondary structure for this domain in the apo state, leading to a slight improvement in the apo prediction (RMSD = 5.97 Å). However, its performance on the holo state diminishes (RMSD = 6.37 Å), reflecting the potential negative effect associated with NMR fine-tuning. This case further illustrates the complementarity between trX2 and trX2 (NMR). In comparison, trX2-D demonstrates superior performance for both states. Through the heuristic sampling and multi-model integration, trX2-D achieves RMSD values of 4.11 Å for the apo state and 4.20 Å for the holo state, which are 33.3% and 6.8% lower than trX2, respectively. Importantly, trX2-D effectively captures both the detailed intra-domain secondary structures and the large-scale inter-domain motion.

### Comparison with other methods

As a comparative benchmark, trX2-D was further evaluated alongside two established methodologies: AlphaFold2^3^ (AF2) and its extension, AF-Cluster^14^, the latter of which produces multiple conformations through clustering of AF2’s MSA input.

To facilitate a thorough understanding of trX2-D’s capabilities, a preliminary comparative analysis between the original trX2 and AF2 was conducted on the apo-holo dataset. As shown in Fig. 2a and Table S4, the original trX2 performs worse than AF2 on both apo and holo states. This disparity can be attributed to two main factors: 1) the relatively lightweight architecture of trX2 compared to AF2; 2) critically, the potential data leakage implied in AF2’s training. The release dates for all 38 apo-holo pairs in our dataset (all prior to 2015; Table S2) predate AF2’s training data cutoff (2020-05), suggesting AF2 might “remember” these native conformations rather than genuinely predicting them. We also find that AF2 tends to favor the holo state, with an average RMSD of 3.32 Å, compared to 4.19 Å for the apo state. For 65.8% of the 38 conformation pairs, the AF2 prediction was closer to the holo state (lower RMSD) than the apo state. This observation is consistent with findings from previous work ^19^. We hypothesize that holo forms are more stable than the apo forms and are therefore more readily predicted by AF2, which excels as a well-trained static structure prediction method.

Regarding AF-Cluster, through the comprehensive MSA clustering, its comprehensive MSA clustering slightly improves apo state predictions over AF2 but offers limited gains for the holo state. The improvements in both states are modest, with RMSD reductions of <0.3 Å (P-value: 0.06 for the apo state and 0.66 for the holo state). In comparison, trX2-D consistently outperforms the original trX2 in predicting both states, achieving RMSD reductions of over 1 Å (P-value: 0.0016 for apo state and 0.0007 for holo state), indicating the effectiveness of our output-based sampling strategy to generate multiple conformations. Consequently, trX2-D exhibited lower RMSD than AF2 for the apo state, and performed comparably to AF-Cluster for both states. For example, as shown in Table S4 and Fig. S7, trX2-D outperforms AF-Cluster for the apo state in 50% (19/38) of targets. It should be noted that trX2-D does not suffer from the data leakage issues implied in AF-Cluster, which further confirms the robustness of its performance.

To gain a deeper understanding of these results, we next assessed the influence of conformational divergence on the performance comparison (Fig. S7b and Fig. S7d). It has been previously observed that AF2 performs poorly for proteins exhibiting significant conformational changes ^19^. For trX2-D, we observe a tendency to outperform AF-Cluster for proteins with large conformational changes. For example, for the targets with RMSD_apo-holo_ over 8.5 Å, trX2-D can generate more accurate structures for both apo and holo states, achieving RMSDs of 7.53 Å and 7.35 Å, respectively, compared to 7.90 Å and 8.64 Å of AF-Cluster. This highlights trX2-D’s better capacity to capture significant structural variability.

To further illustrate these findings with a representative case, we examined an apo-holo pair characterized by a significant conformational change upon calcium binding (Fig. 2d). This pair involves the N-domain of *troponin C*, a crucial regulatory component of the troponin complex that governs muscle contraction. In skeletal and cardiac muscles, *troponin C* binds Ca²⁺, triggering a conformational shift that enables its interaction with *troponin I* and *tropomyosin*, ultimately leading to muscle contraction. The substantial structural divergence observed between the apo (PDB ID: 1SKT^22^) and holo (PDB ID: 1TNQ^23^) states of *troponin C* (RMSD = 7.2 Å, TM-score = 0.54) confirms the conformational change induced by calcium binding.

For this challenging case, both AF2 and AF-Cluster perform better for the holo state (RMSD ∼3 Å) than for the apo state (RMSD ∼4.4 Å). The best-predicted structures from these two methods for the apo and holo states are remarkably similar (TM-score ∼0.95, RMSD ∼2.0 Å), suggesting that neither method successfully captures the conformational change of this protein. In contrast, trX2-D yields a more accurate prediction for the apo state (RMSD=2.87 Å), surpassing both AF2 and AF-Cluster. Although its prediction for the holo state exhibited lower accuracy (RMSD=3.80 Å), the two conformations predicted by trX2-D showed greater structural difference (TM-score = 0.71, RMSD = 2.9 Å) than the AF2-based methods. This reflects trX2-D’s enhanced ability to capture the conformational change.

Another comprehensive dual-conformation dataset originates from Cfold, a recently developed method for predicting alternative protein conformations, which is distinguished by its meticulous training-test split. Specifically, the Cfold dataset was carefully designed to eliminate any conformational redundancy between its training and testing sets. We evaluated our method on 20 targets from this dataset, which exhibit substantial conformational changes and are non-redundant to the training sets of both trX2-D and Cfold. Each target was annotated with “Fold1” and “Fold2” states (see Methods for details on dataset construction and annotation). The information of 20 conformation pairs from Cfold is listed in Table S5.

The results on the Cfold dataset are detailed in Fig. S8 and Table S6. Our method shows enhanced stability, outperforms AF2 specifically for the Fold2 state, and achieves comparable performance to both AF-Cluster and Cfold. Furthermore, trX2-D can capture the large conformational change more effectively than the compared methods (e.g., the case shown in Fig. S9). This independent validation further demonstrates the robustness and broader applicability of our method.

### Application to dynamic structures determined by NMR spectroscopy

The prediction of dynamic protein structures poses a more significant challenge compared to that of proteins exhibiting only two stable conformational states. To evaluate the capabilities of trX2-D in this context, we applied it to a benchmark dataset comprising 62 proteins with dynamic structures solved by NMR spectroscopy. For a balanced comparison, we quantified accuracy for each protein by first determining the minimum RMSD between each predicted conformation and a reference NMR model, and then calculating the mean of these minimum RMSD values across all NMR models. This metric, termed RMSD_avg_, is defined as:

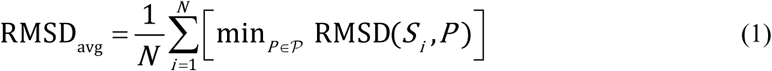

where *N* is the total number of NMR structure models, *S_i_* refers to the *i-*th NMR model, 𝒫 represents the set of predicted conformations, and the minimum RMSD is found by comparing *S_i_* to each prediction *P* within the set 𝒫.

Similar to the above experiments, we compare trX2-D with trX2, AF2, and AF-Cluster on this dynamic protein dataset. As shown in Fig. 5a and Fig. 5b, while all methods face difficulties with this complex task, trX2-D achieves the lowest RMSD_avg_ for most samples, indicating its potential to enhance predictions of dynamic structures. The superior performance of trX2-D over AF-Cluster and AF2 may stem from its greater ability to capture structural diversity, which is essential for accurately representing the native conformational landscape.

**Fig. 5.**
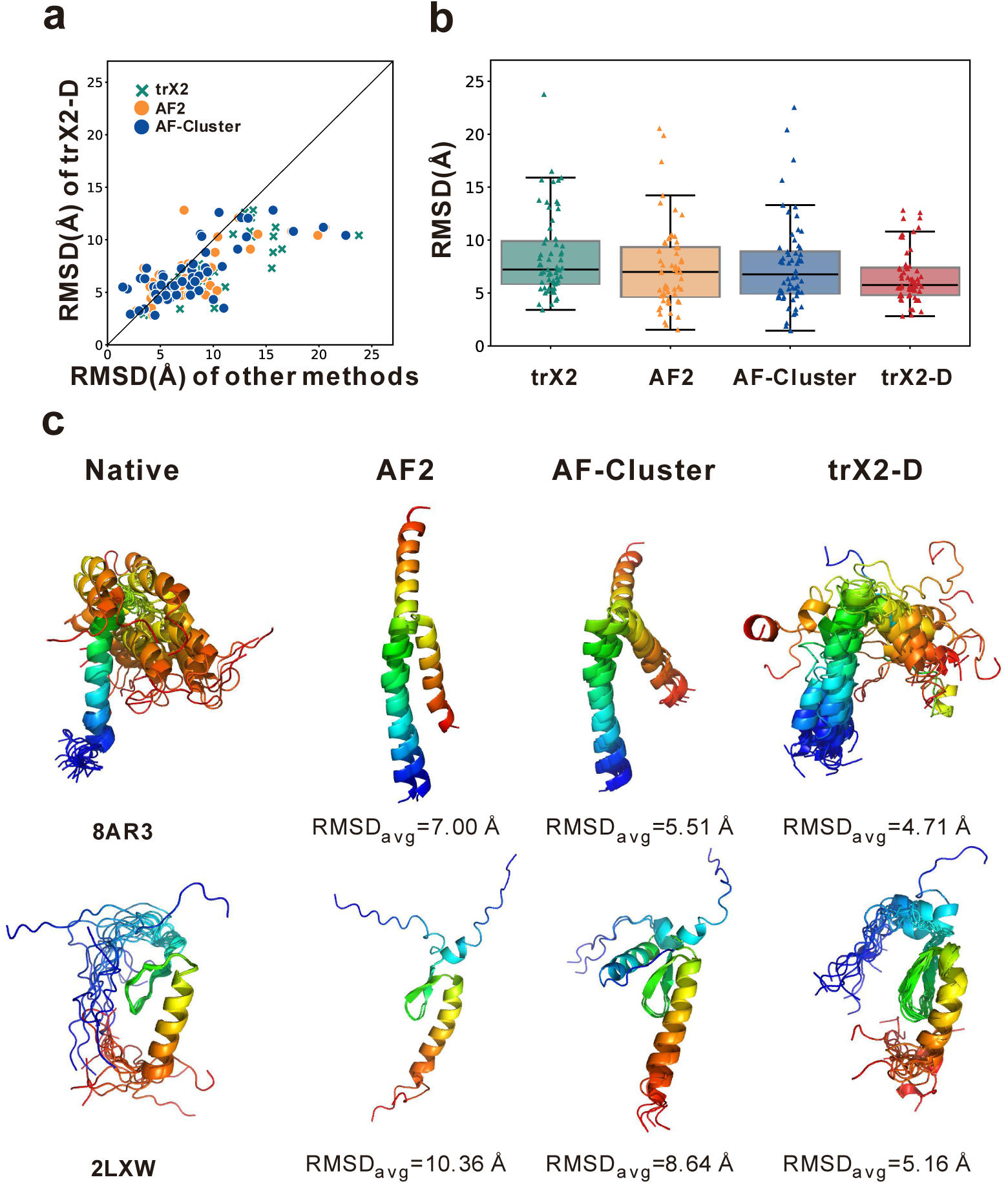
Performance of trX2-D and other methods on dynamic proteins from NMR data. **(a)** head-to-head RMSD comparisons of trX2-D versus trX2, AlphaFold2 (AF2), and AF-Cluster for dynamic proteins; points below the diagonal denote trX2-D superiority. **(b)** boxplot illustrating the RMSD distributions for different methods on the dynamic protein benchmark dataset. **(c)** representative examples of native structures and predictions (PDB IDs: 8AR3, 2LXW). While all methods face challenges, trX2-D demonstrates improved performance in capturing the structural diversity of dynamic proteins.

To illustrate this improvement, we visualize the conformational spaces of the 5 samples exhibiting the largest conformational divergences (Fig. 6). Intriguingly, we observe that these highly divergent cases often involve significant conformational changes within intrinsically disordered regions (IDRs) of the proteins. These disordered regions pose a considerable challenge for AF2 and AF-Cluster, which are primarily trained on stable, well-ordered structures. Consequently, in these challenging cases, predictions from AF2 and AF-Cluster only cover a limited spectrum of the experimental conformational landscapes, resulting in high RMSDs for most conformations and an underestimation of the structural dynamics. In contrast, trX2-D generates a more diverse set of predictions that span a broader conformational range. This enables trX2-D to more effectively capture the structural flexibility inherent in these disordered regions, and consequently achieves lower RMSD_avg_ values than the AF2-based approaches. This observation is reinforced by an analysis of per-residue root mean square fluctuation (RMSF) profiles (Fig. S10). For four of these five highly flexible samples, AF2 and AF-Cluster underestimated structural fluctuations compared to the NMR ensemble, whereas the fluctuations captured by trX2-D models closely correspond with those of the NMR states, reflecting a more faithful depiction of protein flexibility.

**Fig. 6.**
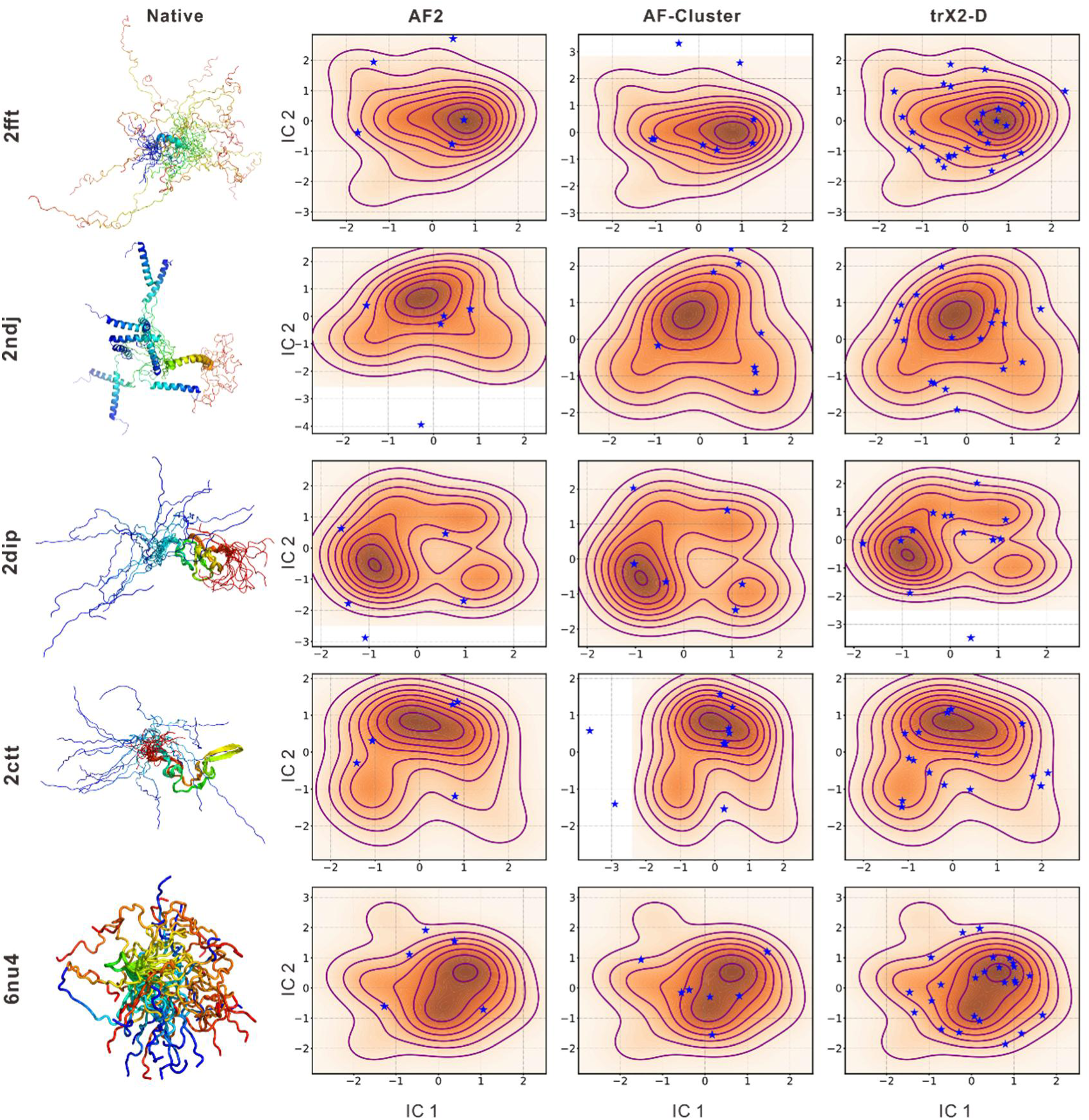
Visualization of conformational space coverage of predicted structures compared to native ensembles for the top 5 proteins with the highest conformational variability. The visualization is based on the first two independent components (IC1 and IC2) derived from Independent Component Analysis (ICA) of Cα coordinates. Density plots show the distribution of NMR models, while the predicted conformations are shown as blue stars. The corresponding NMR ensembles are displayed on the left as rainbow cartoons. These cases highlight the better ability of trX2-D in exploring the conformational space.

Fig. 5c illustrates this trend with two cases characterized by functionally relevant dynamics: the transmembrane and juxtamembrane regions of *Toll-like Receptor 9* (TLR9; PDB ID: 8AR3^24^), where conformational changes drive immune signal transduction; and the *X-linked Inhibitor of Apoptosis Protein* (XIAP)-binding domain of human *X-linked Inhibitor of Apoptosis Factor 1* (XAF1; PDB ID: 2LXW), whose flexibility influences apoptosis regulation. For these targets, both AF2 and AF-Cluster generate predictions with limited conformational spread, resulting in RMSD_avg_ of 7.00 Å/10.36 Å for 8AR3 and 5.51 Å/8.64 Å for 2LXW, respectively. In contrast, trX2-D generates more diverse structural ensembles and achieves lower RMSD_avg_ values of 4.71 Å and 5.16 Å for these two targets, respectively. This improvement highlights the effectiveness of trX2-D in capturing the diversity within these two native ensembles.

Despite these advances, accurately predicting a broader range of dynamic structures remains an open challenge. For example, while trX2-D demonstrates greater prediction diversity than AF2 and AF-Cluster, it still exhibits discrepancies with the native structures (Fig. 5c). These findings highlight the inherent difficulty of modeling dynamic structures and the need for continued methodological improvements. Despite these challenges, trX2-D has shown a distinct and promising capability in capturing structural dynamics.

## Discussion

Structural heterogeneity underpins protein function^25^, yet predicting multiple protein conformational states remains a formidable challenge. Current mainstream methods primarily utilize the multi-conformational information embedded in MSA^10, 14, 15^. In contrast, earlier studies have shown that *de novo*-predicted contact maps obtained through deep learning often contain structural information about multiple states^12^. Motivated by this insight, we observed that the predicted 2D geometries also encode such information, as revealed by their multi-peaked distributions (Fig. S11). However, a systematic approach to utilize 2D geometries for predicting multiple conformations has remained unexplored.

Building upon this observation, we introduce trX2-D, an automated approach for predicting alternative protein conformations by employing a heuristic iterative sampling process on the 2D geometries predicted by trX2. In contrast to existing input-based methods, trX2-D, as an output-driven sampling method, is capable of generating more diverse conformations even without prior knowledge of specific structural states, as its sampling is directly performed on the *de novo*-predicted structural restraints. We have rigorously assessed trX2-D with three independent datasets, including two dual-conformation sets and one dynamic structure set. Benchmarking results demonstrate that trX2-D significantly outperforms trX2 in predicting alternative conformations. On the dual-conformation test sets, trX2-D achieved accuracy comparable to AF-Cluster and Cfold, while outperforming AF2 in predicting alternative states. Unlike MSA clustering methods, trX2-D directly modulates structural geometries, enabling it to capture conformational diversity that was overlooked by other methods (Fig. 2 and Fig. S9). Furthermore, the iterative sampling process holds the potential to predict not just two, but a broader ensemble of functionally relevant conformations. Assessment of the dynamic protein dataset from NMR data reveals that trX2-D can generate structures with greater diversity compared to other methods, underscoring its enhanced capability to model conformational heterogeneity. Importantly, as trX2-D operates solely on the predicted 2D geometries without relying on input MSAs, it also holds potential for broader applications across diverse protein systems (e.g., the orphan proteins that lack sufficient homologous sequences).

Despite these advancements, trX2-D exhibits slightly lower model accuracy than AF-Cluster. This could be partly attributed to the performance difference between the baseline trX2 and AF2 models, along with the potential data leakage in AF2 (Tables S2 and S3). To isolate the contribution of our heuristic iterative strategy from potential biases in training data or baseline model accuracy, we perform a controlled comparison by applying the MSA clustering process to trX2 modeling, i.e., running trX2 using the clustered MSAs from AF-Cluster. The result method, denoted as trX2-Cluster, performs worse than trX2-D for both apo and holo states across the 38 benchmark proteins (Fig. S12). This result confirms the effectiveness of our heuristic iterative strategy over MSA clustering within the trX2 framework.

To further investigate the generalizability of our approach, we explore applying our heuristic iterative strategy to the 2D distance maps predicted by AF2. However, this application failed to generate distinct alternative conformations (AF-HIS in Fig. S13a). This limitation likely arises from the high confidence of AF2’s predicted distance distributions. As a method primarily designed for accurate static structure prediction, the 2D geometries predicted by AF2 are more sharply defined and contain fewer dynamic signals compared to those from trX2, especially the NMR-finetuned version. This assertion is supported by a sharp, unimodal distance distribution observed for AF2 predictions (Fig. S11).

Interestingly, previous research has suggested that AF2 predictions for uniformly subsampled shallow MSAs of size 10 may contain implicit multi-conformational signals^14, 15^. Inspired by this finding, we filter each MSA using HHfilter^26, 27^ to retain only 10 representative sequences. Under these conditions, AF2 predictions exhibit increased diversity in their distance distributions (Fig. S11), consistent with the previous observations. When our heuristic iterative sampling strategy was applied to these modified predictions, it improved the sampling of distinct conformations and enhanced prediction accuracy for alternative conformations (Fig. S13b). However, the overall precision was still limited by the reduced depth of the MSA. These findings highlight a limitation of the sampling strategy, that is, it relies on multi-state information implied in the predicted 2D geometries, thus does not perform well for predictions exhibiting high confidence towards a single, well-defined state (e.g., AF2 predictions using deep MSAs).

Another challenge, which is common for all the existing multi-conformation prediction methods, is the automated and efficient selection of biologically meaningful conformations from the generated ensembles. While trX2-D excels at generating diverse conformational ensembles, it still faces the challenge of selecting representative conformations, especially for dual-conformation proteins. Preliminary efforts using k-means clustering based on standard structural similarity metrics (TM-score, RMSD, and inter-Cα distances^28^) proved insufficient, as illustrated by the 0.2 Å∼0.3 Å higher RMSDs in average after clustering (Fig. S14). This indicates a promising future direction for advancing multi-state structure prediction, that is, exploring more sophisticated conformation selection/clustering strategies to better identify biologically meaningful conformations from generated structures. The incorporation of the physical/biological guidance is a promising approach in this regard.

Conformation generation in trX2-D is primarily powered by energy minimization, which involves both predicted 2D geometries and physical energy terms from Rosetta. Recently, generation models, especially the diffusion models, have shown promise in protein structure generation^29, 30^. However, due to the lack of physical restraints, these generative models alone may struggle to generate conformations that obey the Boltzmann distribution. While several methods (e.g., CONFDIFF^31^, DiG^32^) have made strides in incorporating physical guidance into diffusion and/or sampling procedures, accurately defining force fields and efficiently selecting biologically meaningful conformations continue to be major challenges. The path forward will likely involve a more sophisticated integration of deep generative models with physical/biological restraints, not only to improve the effectiveness of generating diverse conformations but also to better capture those allosteric transitions critical for protein function.

## Methods

### Construction of datasets

#### Test sets

Three benchmark datasets were constructed in this work. The first dataset consists of 38 apo-holo protein pairs collected from a recent work by Saldano *et al*^19^. From their original set of 91 pairs, 87 with identical sequences between the apo and holo states were initially selected. To focus on pairs with substantial conformational differences, we only retained the 38 protein pairs with significant conformational change, defined as having a TM-score_apo-holo_ value below 0.8 or an RMSD_apo-holo_ value above 6 Å. While a TM-score_apo-holo_ of 0.8 is frequently used as the cutoff to identify significant conformational changes^10, 12, 16^, our analysis revealed cases where high TM-scores between states can also coincide with substantial structural differences, as indicated by RMSD_apo-holo_ values exceeding 6 Å (Fig. S9 and Fig. S15). For example, for the DAHPS enzyme, the transition between states involves significant interdomain variation (apo PDB ID: 1RZM, holo PDB ID: 1VR6; Fig. S15a). However, its TM-score_apo-holo_ value is over the 0.8 cutoff. Therefore, to ensure a robust evaluation, we incorporated such cases into our benchmark set.

The second dataset was obtained from the Cfold benchmark set^10^, which includes 243 dual-conformation proteins with pairwise TM-score<0.8. For consistency, we only considered 155 samples for which Cfold provided the predicted structures in its Zenodo repository (https://zenodo.org/records/10837082). To prevent data leakage, we further filtered this dataset using the cd-hit^33, 34^ (V4.8.1) program at a 40% sequence identity threshold relative to our training set. After this step, we removed samples with sequence lengths >300 to save computational time, resulting in 20 unique samples. For conformation annotation, we categorized proteins based on the RMSD of their AF2 predictions. For each conformation pair, the conformation with lower RMSD in at least 3 out of 5 AF2 predictions was labeled as “Fold1”, while the other conformation in the pair was designated as “Fold2”^16^.

The third dataset is derived from the 292 dynamic proteins identified by NMR spectroscopy reserved from the dataset used to fine-tune trX2 (see below), each protein with an average of 19 NMR models. The proteins sharing over 30% sequence identity relative to all the training sets were excluded, resulting in 118 samples. To ensure sufficient conformational diversity within each sample, only proteins with conformation ensembles among which the minimum pairwise TM-score below 0.8 or the maximum pairwise RMSD over 6 Å were retained, resulting in 92 proteins. Subsequently, 29 proteins were excluded due to insufficient MSA depth (<10) for running AF-Cluster. Furthermore, one protein (PDB ID: 6xry) with extreme conformational dynamics (maximum pairwise RMSD = 33.9 Å) was also excluded, as all evaluated methods failed to generate structure ensembles with RMSD_avg_ below 26 Å for this target. The final dataset thus consists of 62 proteins with dynamic structures.

Duplicate samples across the above three datasets were removed to eliminate redundancy. The final datasets consist of 38 apo-holo proteins, 20 two-state proteins, and 62 dynamic proteins, respectively.

#### X-ray training set

This training set was derived from the 15,051 X-ray protein chains collected in trRosetta^5^. These proteins are non-redundant (sequence identity <30%), released before 2018-05-01 in the PDB database, resolved by X-ray crystallography, and have pre-constructed MSAs with at least 100 homologous sequences. To prevent data leakage during benchmark evaluations, any training set chains sharing >50% sequence identity with proteins in the benchmark test set were removed, resulting in a final training set of 14,275 chains.

#### NMR training set

This training set was derived from a dataset of 8,038 monomeric proteins with experimentally determined dynamic structures from NMR spectroscopy. A two-stage filtering procedure was employed to ensure non-redundancy and prevent data leakage. First, the chains were clustered at 60% sequence identity using CD-HIT. 95% of the resulting 4,746 clusters (7,454 chains) were randomly selected for the initial training set, while the remaining 5% of clusters (292 chains) were reserved for the test set construction. Second, to further prevent data leakage during benchmark evaluations, any training chains sharing >50% sequence identity with proteins in the benchmark test set were removed, resulting in a final training set of 7,269 chains.

### Experimental Setup

#### MSA Generation

MSAs for all proteins were generated using MMseqs2^35^ (v13.45111) by searching the UniRef50 database (E-value threshold: 0.001; maximum 20,000 target sequences per query). Unless otherwise specified, these MSAs served as the common input for all structure prediction methods to ensure fair comparison.

#### Structure Prediction Methods

We compared trX2-D to the following structure prediction methods:

#### trRosettaX2 (trX2)

The trX2 protocol was used to predict 2D geometric constraints from the input MSA, which guided structure folding via energy minimization in Rosetta. To sample diverse conformations, rather than relying on a fixed end-to-end prediction, we executed 50 independent energy minimization processes for each target. These processes incorporated randomness in both their initialization and optimization steps, ultimately yielding an ensemble of 50 structures.

#### AlphaFold2 (AF2)

The standard AlphaFold2 without structural templates was used to generate predictions. For each target, the top 5 models ranked by the predicted Local Distance Difference Test (pLDDT) scores were selected for evaluation.

#### AF-Cluster

Following the AF-Cluster pipeline (https://github.com/HWaymentSteele/ <u>AF_Cluster</u>), each target’s MSA was clustered into 20 sub-MSAs: ten containing 10 sequences each and ten containing 100 sequences each. AlphaFold2 (without templates) was run on each sub-MSA individually, producing one structure per sub-MSA and thus a total ensemble of 20 structures per target.

#### Cfold

The Cfold predictions were obtained directly from the published dataset on the Zenodo repository (https://zenodo.org/records/10837082). We specifically used structures generated by Cfold’s MSA clustering strategy, which was reported to outperform the alternative dropout strategy in the original study.

#### Evaluation Metric

The quality of the predicted models was evaluated by RMSD. For each experimentally determined conformation of a target, RMSD values comparing all generated structures against this conformation were computed utilizing the TM-score^36^ program. Then the minimum RMSD value was selected to represent the accuracy for that specific conformation.

### NMR fine-tuning of trRosettaX2

To improve the ability to capture the conformational changes, we fine-tuned the pretrained trX2 (described in Text S1) on the dynamic structures from the NMR training set. The loss function was adapted to consider all the conformations of each sample. Specifically, for each protein, we computed the loss function between the predicted structure and all native conformations and selected the minimum loss for backpropagation. This process can be written as:

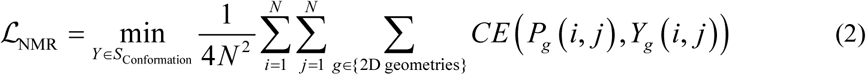

where *S*_Conformation_ refers to the set of native conformations; *N* is the number of residues; *CE*() is the cross-entropy function; *P_g_*(*i*,*j*) is the predicted probability distribution for the 2D geometry *g* between residues *i* and *j*; *Y_g_*(*i*,*j*) is the corresponding ground truth one-hot encoding derived from the native conformation *Y*.

Throughout the training, we used the Adam optimizer with a learning rate of 0.0001 to minimize the loss function ℒ_NMR_.

### Heuristic iterative process for trX2-D

The heuristic iterative process in trX2-D is designed to generate a diverse ensemble of protein conformations from the 2D geometries predicted by trX2 and trX2 (NMR). At each iteration, geometry information from the prior iteration’s 3D conformation is selectively excluded from the current 2D geometries. These updated 2D geometries are subsequently used to generate a new conformation through energy minimization. Sufficient iterations of this process can yield a diverse conformational set. This procedure is illustrated in Figs. 1C and S2.

Formally, let 𝒢_*n*_ denote the set of 2D geometries (1 distance + 3 orientations) at the *n*-th iteration. *S*_*n*_ is the corresponding 3D structure generated through energy minimization based on 𝒢_*n*_. The *n*-th iteration aims to update 𝒢_*n*_ and *S*_*n*_ to 𝒢_*n*+1_ and *S*_*n*+1_, respectively. Once the iteration process terminates, all generated structures, {*S*_*n*_}, are collected to form the predicted ensemble of conformations.

For convenience, let *p*_*n*_ ∈ 𝒢_*n*_ represent the probability distribution of one of four defined geometries for a specific pair of residues. A sharp and unimodal *p*_*n*_ signifies a highly stable geometric relationship between the corresponding residue pair, while a broad or bimodal *p*_*n*_ may imply variability for this residue pair. Based on this hypothesis, we design a decay-and-smooth procedure to update *p*_*n*_ to *p*_*n*+1_ (Fig. S2), which is detailed as follows (‖⋅‖ refers to the *L* norm):

1. if ‖*p*_*n*_‖ < 0.5, indicating potential conformational variation at this residue pair, *p*_*n*_ will be decayed based on the geometry value calculated for the corresponding residue pair in the 3D structure *S*_*n*_. This operation is intended to remove the information inherent in the previously generated 3D conformation and to focus on the alternative conformation information implied in the remaining distribution regions. The decayed distribution is then normalized and smoothed with a Gaussian filter to ensure structural regularity during energy minimization (see Fig. S16 for an example).
2. if ‖*p*_*n*_‖ ≥ 0.5, indicating this residue pair is highly stable, *p*_*n*_ will remain unchanged. In total, the update rule is defined as:

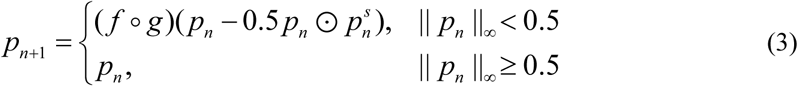

where ⊙ denotes element-wise multiplication, *g* represents normalization, and *f* denotes Gaussian smoothing. 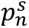 refers to the distribution (i.e., one-hot coding) calculated from the 3D structure *S_n_*.

The updated 2D geometries 𝒢_*n*+1_ are obtained by updating all four types of geometry across all residue pairs in the protein. These updated geometries are then used to generate a new conformation *S*_*n*+1_. The iterative process terminates when the distributions for all residue pairs have converged (change < 0.01).

## Availability

The web server of trX2 is available at: https://yanglab.qd.sdu.edu.cn/trRosetta/. The source codes for trX2-D are available at: https://github.com/YangLab-SDU/trRosettaX2-Dynamics/. The three benchmark datasets for trX2-D are available at https://yanglab.qd.sdu.edu.cn/trRosetta/benchmark_dynamics/.

## Acknowledgements

This work is supported by the following funding sources: National Key Research and Development Program of China (2024YFA0916901), National Natural Science Foundation of China (NSFC T2225007, T2222012, 32430063), Postdoctoral Fellowship Program and China Postdoctoral Science Foundation (BX20240212), and Fundamental Research Funds for the Central Universities.

