## Supplementary Materials for "Generating dynamic structures through physics-based sampling of trRosettaX2-predicted inter-residue geometries"

**Supplement Text 1. Performance of trRosettaX2 on protein static structure prediction**

**Architecture of trRosettaX2.** The trRosettaX2 architecture is primarily inspired by AlphaFold2 (AF2), but several modifications were made to optimize performance under limited computational resources, which is summarized in Table S1. First, we introduced an MSA-based language model ESM-MSA ^1^, which was pretrained on the MSAs of UniRef50 sequences, to generate an initial MSA embedding and 2D attention maps. Second, to accommodate the constraints of GPU memory (A100 40GB), we utilize Direct Coupling Analysis (DCA) scores for embedding the "extra MSA" rather than relying on transformer blocks. Third, observing that the pure transformer network struggles to converge with a "batch size=1" configuration, we introduced the multi-scale convolutional neural network Res2Net (R2N) ^2^ to assist attention-based triangle updates in the Evoformer. This modified module, termed trFormer (Fig. S1a), combines the strengths of transformers and convolutional networks—leveraging the former for global topology modeling and the latter for capturing local details. Notably, we only use 12 trFormer blocks, a significant reduction from the 48 Evoformer blocks used in AlphaFold2.

During inference, the input MSA is transformed into two representations: the MSA representation (i.e., the MSA embedding generated by ESM-MSA) and the pair representation (including the direct couplings derived from MSA and the attention maps from ESM-MSA). These two representations are then iteratively updated by 12 trFormer blocks. The first row of the updated MSA representation, along with the full pair representation, is fed into the structure module to predict the full-atom 3D structures.

The loss function of trRosettaX2 consists of several independent items, including the frame-aligned-point-error (FAPE) loss $\mathcal{L}_{FAPE}$, the 2D geometries loss $\mathcal{L}_{2D}$, the torsion loss $\mathcal{L}_{torsion}$, the pLDDT loss $\mathcal{L}_{pLDDT}$, the clash loss $\mathcal{L}_{clash}$ , and the bond loss $\mathcal{L}_{bond}$. In total, the loss function can be written as:

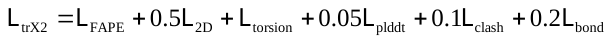
 (S1)

The training of trRosettaX2 is performed on a single A100 40GB GPU and takes around 10 days, using the training set consisting of 14,275 protein crystal structures introduced in Methods.

**Performance of trRosettaX2 for static structure prediction.** To validate the effectiveness of the above modifications, we trained and evaluated several model variants on 91 domains from the CASP14 experiments. The evaluation is based on the average TM-score on the CASP14 dataset, using trRosettaX_FM (trX_FM; the template-free version of trRosettaX), RoseTTAFold (RF; the pyRosetta version), and AlphaFold2 (AF2) as reference models. During prediction, all models use the same MSA as input and do not rely on structural templates.

According to Fig. S1b, directly adopting the Evoformer architecture performs poorly, with an average TM-score of 0.576 on the CASP14 domains, which is even lower than the Res2Net-based trX_FM. This reflects the incompatibility between pure transformer networks and the training configuration we employed under limited resources (e.g., batch size of 1 and only 12 blocks). However, by incorporating Res2Net with the transformer (i.e., trFormer), the model's average TM-score significantly improves by 27.5%, surpassing trX_FM by 11.6%. Training with binary-sub-sampling^3^ further increases the TM-score by approximately 2.5%, outperforming RF. With the introduction of ESM-MSA representations, the TM-score improves again by about 4%. Finally, using the same input (i.e., MSA), trX2 achieved an average TM-score of 0.781, 18.7% higher than trX_FM and 5.8% higher accuracy than RF. Additionally, the trX2 performance is competitive with AF2 (TM-score 0.781 vs 0.843), despite using fewer parameters (0.3B vs 0.9B) and computational resources (1 A100 GPU vs 128 TPUv3 cores). These results illustrate the effectiveness of the modification to accommodate the limited computational resources. On the 94 domains from CASP15, we can also observe a similar comparison (Fig. S1c). trX2 represents a good trade-off of modeling accuracy and parameters/training cost compared to other state-of-the-art methods, further demonstrating its efficiency as a static structure prediction method.

**Table S1 |** Methodological comparison between trX2, AF2, and RF.

| **Methods** | **Approach** | **Neural network architecture** | **Number of parameters** | **Training resources** | **Training time** |
| --- | --- | --- | --- | --- | --- |
| trRosettaX2 | End-to-end/two-steps | R2N-enhanced transformer  (trFormer,  12 blocks) | 30 M | 1 A100 40GB GPU | ~ 10 days |
| AlphaFold2 | End-to-end | Evoformer  (48 blocks)  +structure module  (8 blocks) | 93 M | 128 TPU v3 cores | ~ 2 weeks |
| RoseTTAFold (pyRosetta) | Two-steps | 3-track transformer  (13 blocks) | 130 M | 8 V100 32GB GPUs | ~ 4 weeks |
| RoseTTAFold (e2e) | End-to-end | 3-track transformer  (13 blocks)  +SE3-transformer  (2 layers) | 137 M |  |  |

**Table S2 |** Information of 38 paired apo and holo proteins, along with RMSD and TM-score between apo and holo states.

| **Apo state** | | **Holo state** | | **RMSD(Å)** | **TM-score** |
| --- | --- | --- | --- | --- | --- |
| **PDB ID** | **Release Date** | **PDB ID** | **Release Date** | **between apo and holo** | **between apo and holo** |
| 2LAO | 1993-02-25 | 1LAH | 1993-10-06 | 4.75 | 0.70 |
| 1NTR | 1994-09-16 | 1KRX | 2002-01-10 | 4.55 | 0.63 |
| 1CFC | 1995-08-02 | 5DOW | 2015-09-11 | 9.25 | 0.38 |
| 1MUT | 1995-09-14 | 1PUN | 2003-06-25 | 4.98 | 0.58 |
| 1LIP | 1995-09-21 | 1JTB | 1996-12-03 | 3.29 | 0.66 |
| 4AKE | 1995-12-29 | 2ECK | 1996-12-16 | 6.95 | 0.69 |
| 1SYM | 1996-05-29 | 1XYD | 2004-11-09 | 6.69 | 0.54 |
| 1AEL | 1996-07-30 | 1URE | 1996-06-17 | 2.69 | 0.77 |
| 2CJO | 1997-02-06 | 1ROE | 1995-11-24 | 4.55 | 0.58 |
| 2RCS | 1997-05-14 | 1AJ7 | 1997-05-15 | 7.21 | 0.59 |
| 1URP | 1998-04-03 | 2DRI | 1994-09-23 | 4.20 | 0.72 |
| 1SKT | 1998-04-06 | 1TNQ | 1995-07-07 | 7.19 | 0.54 |
| 1EX6 | 2000-05-01 | 1EX7 | 2000-05-01 | 4.37 | 0.78 |
| 1F3Y | 2000-06-06 | 1JKN | 2001-07-12 | 5.08 | 0.79 |
| 1FMF | 2000-08-17 | 1ID8 | 2001-04-04 | 4.74 | 0.61 |
| 1GH1 | 2000-10-29 | 1CZ2 | 1999-09-01 | 3.04 | 0.68 |
| 1IJA | 2001-04-25 | 2KID | 2009-05-01 | 4.54 | 0.79 |
| 1JFJ | 2001-06-20 | 1JFK | 2001-06-21 | 14.73 | 0.47 |
| 1K2H | 2001-09-27 | 1ZFS | 2005-04-20 | 6.59 | 0.49 |
| 1GUD | 2002-01-24 | 1RPJ | 1999-02-04 | 4.45 | 0.72 |
| 1MO7 | 2002-09-08 | 1MO8 | 2002-09-08 | 4.93 | 0.79 |
| 1ORM | 2003-03-14 | 1QJ8 | 1999-06-23 | 6.13 | 0.51 |
| 1WD7 | 2004-05-12 | 1WCW | 2004-05-06 | 4.12 | 0.71 |
| 1W4U | 2004-07-29 | 1UR6 | 2003-10-27 | 3.01 | 0.75 |
| 1VR6 | 2005-02-14 | 1RZM | 2003-12-24 | 10.10 | 0.81 |
| 1Z15 | 2005-03-03 | 1Z17 | 2005-03-03 | 6.45 | 0.66 |
| 2CG7 | 2006-02-27 | 2CG6 | 2006-02-27 | 6.25 | 0.55 |
| 2NLN | 2006-10-20 | 1RRO | 1992-08-27 | 3.90 | 0.63 |
| 2UZ5 | 2007-04-25 | 2VCD | 2007-09-20 | 3.37 | 0.78 |
| 2JU3 | 2007-08-14 | 2JU8 | 2007-08-15 | 3.17 | 0.73 |
| 2JWW | 2007-10-25 | 1RTP | 1993-05-14 | 2.79 | 0.73 |
| 2K43 | 2008-05-28 | 2K8R | 2008-09-22 | 3.93 | 0.73 |
| 2KQ2 | 2009-10-24 | 2KW4 | 2010-03-31 | 14.14 | 0.28 |
| 2KXL | 2010-05-10 | 2K0G | 2008-02-02 | 6.09 | 0.66 |
| 2L50 | 2010-10-22 | 2L51 | 2010-10-22 | 4.56 | 0.73 |
| 2LKC | 2011-10-10 | 2LKD | 2011-10-10 | 7.47 | 0.72 |
| 4LP5 | 2013-07-15 | 4P2Y | 2014-03-05 | 11.69 | 0.74 |
| 6OY9 | 2019-05-14 | 1A0R | 1997-12-05 | 3.94 | 0.75 |

**Table S3 |** Impact of the different components on the structure modeling accuracy on 38 apo and holo states proteins.

| **Component** | **RMSD against apo state (Å)** | **RMSD against holo state (Å)** |
| --- | --- | --- |
| 1) baseline (trX2) | 5.40 | 5.16 |
| 2) baseline + NMR training set | 5.13 | 4.38 |
| 3) baseline + iteration | 4.90 | 4.53 |
| 4) baseline + NMR training set + iteration | 4.71 | 4.00 |
| 5) integration of (1) and (2) | 4.57 | 4.07 |
| 6) integration of (3) and (4) (trX2-D) | 4.18 | 3.60 |

**Table S4 |** Comparison of AF2, AF-Cluster, trX2, and trX2-D on apo and holo states proteins

| **Apo state** | **RMSD against apo state(Å)** | | | | **Holo state** | **RMSD against holo state(Å)** | | | |
| --- | --- | --- | --- | --- | --- | --- | --- | --- | --- |
| **PDB ID** | **AF2** | **AF- Cluster** | **trX2** | **trX2-D** | **PDB ID** | **AF2** | **AF- Cluster** | **trX2** | **trX2-D** |
| 2LAO | 4.62 | 2.92 | 2.91 | 2.83 | 1LAH | 0.41 | 0.49 | 3.00 | 1.59 |
| 1NTR | 4.17 | 4.32 | 4.40 | 4.10 | 1KRX | 3.27 | 3.44 | 3.22 | 2.92 |
| 1CFC | 5.51 | 5.92 | 7.61 | 5.78 | 5DOW | 6.06 | 5.62 | 8.41 | 6.72 |
| 1MUT | 3.70 | 3.71 | 3.96 | 3.75 | 1PUN | 3.78 | 4.09 | 4.26 | 3.90 |
| 1LIP | 1.58 | 1.64 | 2.09 | 1.88 | 1JTB | 2.92 | 2.83 | 2.82 | 2.63 |
| 4AKE | 6.07 | 2.69 | 3.59 | 3.25 | 2ECK | 0.93 | 1.70 | 5.10 | 1.79 |
| 1SYM | 5.50 | 3.39 | 5.06 | 4.94 | 1XYD | 2.01 | 2.13 | 3.48 | 3.43 |
| 1AEL | 2.51 | 2.46 | 2.77 | 2.49 | 1URE | 1.50 | 1.49 | 1.76 | 1.71 |
| 2CJO | 1.29 | 1.43 | 1.72 | 1.34 | 1ROE | 4.33 | 4.34 | 4.26 | 4.21 |
| 2RCS | 5.79 | 4.07 | 7.57 | 3.43 | 1AJ7 | 1.96 | 2.69 | 9.50 | 4.60 |
| 1URP | 3.79 | 0.82 | 2.75 | 2.47 | 2DRI | 0.52 | 0.50 | 2.33 | 1.12 |
| 1SKT | 4.46 | 4.41 | 3.93 | 2.87 | 1TNQ | 3.00 | 3.18 | 4.00 | 3.80 |
| 1EX6 | 0.93 | 1.51 | 3.10 | 3.00 | 1EX7 | 1.11 | 0.94 | 2.07 | 1.63 |
| 1F3Y | 3.13 | 3.71 | 4.32 | 3.80 | 1JKN | 2.59 | 2.18 | 2.68 | 2.47 |
| 1FMF | 1.83 | 1.89 | 2.49 | 2.19 | 1ID8 | 4.57 | 4.43 | 4.40 | 4.15 |
| 1GH1 | 2.38 | 2.09 | 2.18 | 1.99 | 1CZ2 | 2.67 | 2.59 | 2.53 | 2.48 |
| 1IJA | 3.23 | 3.42 | 5.92 | 6.18 | 2KID | 2.04 | 2.10 | 6.45 | 4.76 |
| 1JFJ | 13.54 | 13.57 | 10.32 | 10.23 | 1JFK | 12.91 | 12.92 | 11.52 | 6.38 |
| 1K2H | 6.49 | 5.55 | 5.32 | 5.37 | 1ZFS | 2.14 | 2.16 | 3.35 | 3.14 |
| 1GUD | 3.74 | 2.61 | 3.58 | 3.35 | 1RPJ | 0.97 | 1.14 | 2.90 | 2.77 |
| 1MO7 | 3.80 | 4.05 | 5.51 | 4.23 | 1MO8 | 3.43 | 3.35 | 4.67 | 4.04 |
| 1ORM | 5.71 | 6.05 | 5.26 | 4.82 | 1QJ8 | 0.95 | 0.67 | 2.10 | 1.49 |
| 1WD7 | 3.22 | 4.64 | 14.24 | 4.33 | 1WCW | 0.87 | 1.06 | 14.20 | 1.52 |
| 1W4U | 2.26 | 2.20 | 2.26 | 2.04 | 1UR6 | 2.16 | 2.16 | 2.46 | 2.02 |
| 1VR6 | 1.56 | 1.56 | 6.17 | 4.11 | 1RZM | 9.21 | 5.09 | 4.51 | 4.2 |
| 1Z15 | 6.10 | 5.25 | 4.74 | 4.50 | 1Z17 | 0.65 | 0.66 | 2.64 | 1.63 |
| 2CG7 | 0.99 | 1.26 | 6.45 | 2.16 | 2CG6 | 6.22 | 3.84 | 9.57 | 2.52 |
| 2NLN | 3.92 | 3.88 | 3.61 | 3.43 | 1RRO | 0.64 | 0.61 | 1.11 | 1.13 |
| 2UZ5 | 3.01 | 3.09 | 3.10 | 2.96 | 2VCD | 1.38 | 1.48 | 1.86 | 1.50 |
| 2JU3 | 2.70 | 2.71 | 2.68 | 2.52 | 2JU8 | 1.79 | 1.73 | 1.71 | 1.71 |
| 2JWW | 2.63 | 2.61 | 2.85 | 2.82 | 1RTP | 0.38 | 0.47 | 1.42 | 1.45 |
| 2K43 | 2.77 | 2.85 | 2.80 | 2.67 | 2K8R | 4.01 | 3.80 | 3.68 | 3.39 |
| 2KQ2 | 8.97 | 9.43 | 9.67 | 8.37 | 2KW4 | 10.79 | 9.10 | 9.13 | 8.54 |
| 2KXL | 5.09 | 2.72 | 3.23 | 2.74 | 2K0G | 3.38 | 2.96 | 3.88 | 3.23 |
| 2L50 | 3.10 | 3.17 | 10.13 | 5.10 | 2L51 | 3.37 | 3.33 | 9.88 | 4.70 |
| 2LKC | 7.39 | 7.72 | 7.64 | 6.01 | 2LKD | 4.10 | 4.43 | 5.05 | 4.56 |
| 4LP5 | 9.88 | 9.04 | 8.94 | 9.18 | 4P2Y | 8.98 | 10.48 | 10.53 | 10.94 |
| 6OY9 | 1.83 | 2.48 | 20.20 | 11.50 | 1A0R | 4.17 | 4.58 | 19.78 | 11.88 |
| **Average** | **4.19** | **3.86** | **5.40** | **4.18** | **Average** | **3.32** | **3.18** | **5.16** | **3.60** |

**Table** **S5 |** Information of 20 paired dual-conformation proteins from the Cfold benchmark set, the definitions of Fold1 and Fold2 are provided by Chakravarty D, *et al* ^4^.

| **Fold1 state** | | **Fold2 state** | | **RMSD(Å)** | **TM-score** |
| --- | --- | --- | --- | --- | --- |
| **PDB ID** | **Release Date** | **PDB ID** | **Release Date** | **between Fold1 and Fold2** | **between Fold1 and Fold2** |
| 1RLM | 2003-11-26 | 2HF2 | 2006-06-22 | 3.58 | 0.79 |
| 2BBW | 2005-10-17 | 2AR7 | 2005-08-19 | 5.16 | 0.75 |
| 2IEY | 2006-09-19 | 2IEZ | 2006-09-19 | 5.49 | 0.70 |
| 2NXE | 2006-11-17 | 2ZBP | 2007-10-26 | 11.75 | 0.78 |
| 2V9R | 2007-08-25 | 2V9Q | 2007-08-25 | 3.87 | 0.67 |
| 3BIS | 2007-11-30 | 3BIK | 2007-11-30 | 3.86 | 0.73 |
| 3PIL | 2010-11-07 | 3PIM | 2010-11-07 | 7.39 | 0.74 |
| 3T9L | 2011-08-03 | 4A3P | 2011-10-03 | 4.92 | 0.61 |
| 3UV5 | 2011-11-29 | 7L6X | 2020-12-24 | 11.97 | 0.57 |
| 5JGK | 2016-04-20 | 5JGL | 2016-04-20 | 6.63 | 0.68 |
| 5O37 | 2017-05-23 | 5O2K | 2017-05-21 | 6.81 | 0.58 |
| 5OVZ | 2017-08-30 | 4P0I | 2014-02-21 | 4.58 | 0.67 |
| 6DZX | 2018-07-05 | 6CVA | 2018-03-27 | 3.91 | 0.73 |
| 6HNI | 2018-09-15 | 6HNK | 2018-09-15 | 4.31 | 0.75 |
| 6P8R | 2019-06-07 | 6P8O | 2019-06-07 | 6.15 | 0.58 |
| 6SVF | 2019-09-18 | 6GGP | 2018-05-03 | 5.68 | 0.58 |
| 6UHI | 2019-09-27 | 6UHS | 2019-09-27 | 3.76 | 0.68 |
| 6ZJB | 2020-06-28 | 6ZJD | 2020-06-28 | 6.74 | 0.76 |
| 7RDT | 2021-07-11 | 7RDS | 2021-07-11 | 8.75 | 0.51 |
| 8DP6 | 2022-07-15 | 8DP7 | 2022-07-15 | 4.40 | 0.72 |

**Table S6** **|** Comparison of AF2, AF-Cluster, Cfold, and trX2-D on 20 dual-conformation proteins

| **Fold1 state** | **RMSD against Fold1 state(Å)** | | | | **Fold2 state** | **RMSD against Fold2 state(Å)** | | | |
| --- | --- | --- | --- | --- | --- | --- | --- | --- | --- |
| **PDB ID** | **AF2** | **AF- Cluster** | **Cfold** | **trX2-D** | **PDB ID** | **AF2** | **AF- Cluster** | **Cfold** | **trX2-D** |
| 1RLM | 0.52 | 0.73 | 0.99 | 1.47 | 2HF2 | 3.34 | 2.36 | 3.12 | 2.93 |
| 2BBW | 2.48 | 2.13 | 3.50 | 2.56 | 2AR7 | 4.91 | 3.64 | 2.57 | 3.97 |
| 2IEY | 15.91 | 15.88 | 15.74 | 15.48 | 2IEZ | 20.18 | 20.17 | 20.10 | 18.72 |
| 2NXE | 5.79 | 5.00 | 5.78 | 4.50 | 2ZBP | 10.55 | 10.24 | 6.40 | 3.77 |
| 2V9R | 0.78 | 0.88 | 2.17 | 2.22 | 2V9Q | 3.33 | 2.67 | 6.01 | 3.67 |
| 3BIS | 2.24 | 2.42 | 3.97 | 2.54 | 3BIK | 2.73 | 2.02 | 2.27 | 3.07 |
| 3PIL | 1.16 | 1.28 | 1.69 | 2.73 | 3PIM | 7.93 | 7.91 | 6.93 | 7.59 |
| 3T9L | 1.81 | 1.86 | 3.11 | 3.25 | 4A3P | 3.93 | 1.96 | 1.56 | 3.55 |
| 3UV5 | 6.88 | 6.21 | 11.89 | 3.71 | 7L6X | 5.07 | 5.62 | 1.43 | 3.40 |
| 5JGK | 0.96 | 1.18 | 1.41 | 2.57 | 5JGL | 6.19 | 5.84 | 5.98 | 6.65 |
| 5O37 | 1.04 | 0.96 | 1.33 | 1.69 | 5O2K | 5.56 | 4.68 | 5.20 | 3.37 |
| 5OVZ | 0.50 | 0.55 | 1.45 | 1.82 | 4P0I | 4.48 | 1.94 | 3.05 | 3.47 |
| 6DZX | 1.70 | 1.73 | 1.21 | 2.34 | 6CVA | 3.82 | 3.32 | 2.94 | 3.63 |
| 6HNI | 0.64 | 0.51 | 1.13 | 1.65 | 6HNK | 4.66 | 1.57 | 3.76 | 3.74 |
| 6P8R | 3.73 | 3.88 | 1.67 | 8.43 | 6P8O | 5.89 | 5.83 | 4.41 | 4.58 |
| 6SVF | 0.37 | 0.38 | 1.70 | 1.35 | 6GGP | 5.68 | 5.04 | 4.93 | 5.12 |
| 6UHI | 1.32 | 1.69 | 3.76 | 2.48 | 6UHS | 2.36 | 2.07 | 3.31 | 2.59 |
| 6ZJB | 1.32 | 2.00 | 4.91 | 1.53 | 6ZJD | 4.61 | 3.70 | 1.91 | 3.42 |
| 7RDT | 1.84 | 1.83 | 1.77 | 2.32 | 7RDS | 8.97 | 8.71 | 8.59 | 8.08 |
| 8DP6 | 0.69 | 0.78 | 1.54 | 1.99 | 8DP7 | 1.45 | 2.30 | 1.60 | 2.20 |
| **Average** | **2.58** | **2.59** | **3.54** | **3.33** | **Average** | **5.78** | **5.08** | **4.80** | **4.87** |

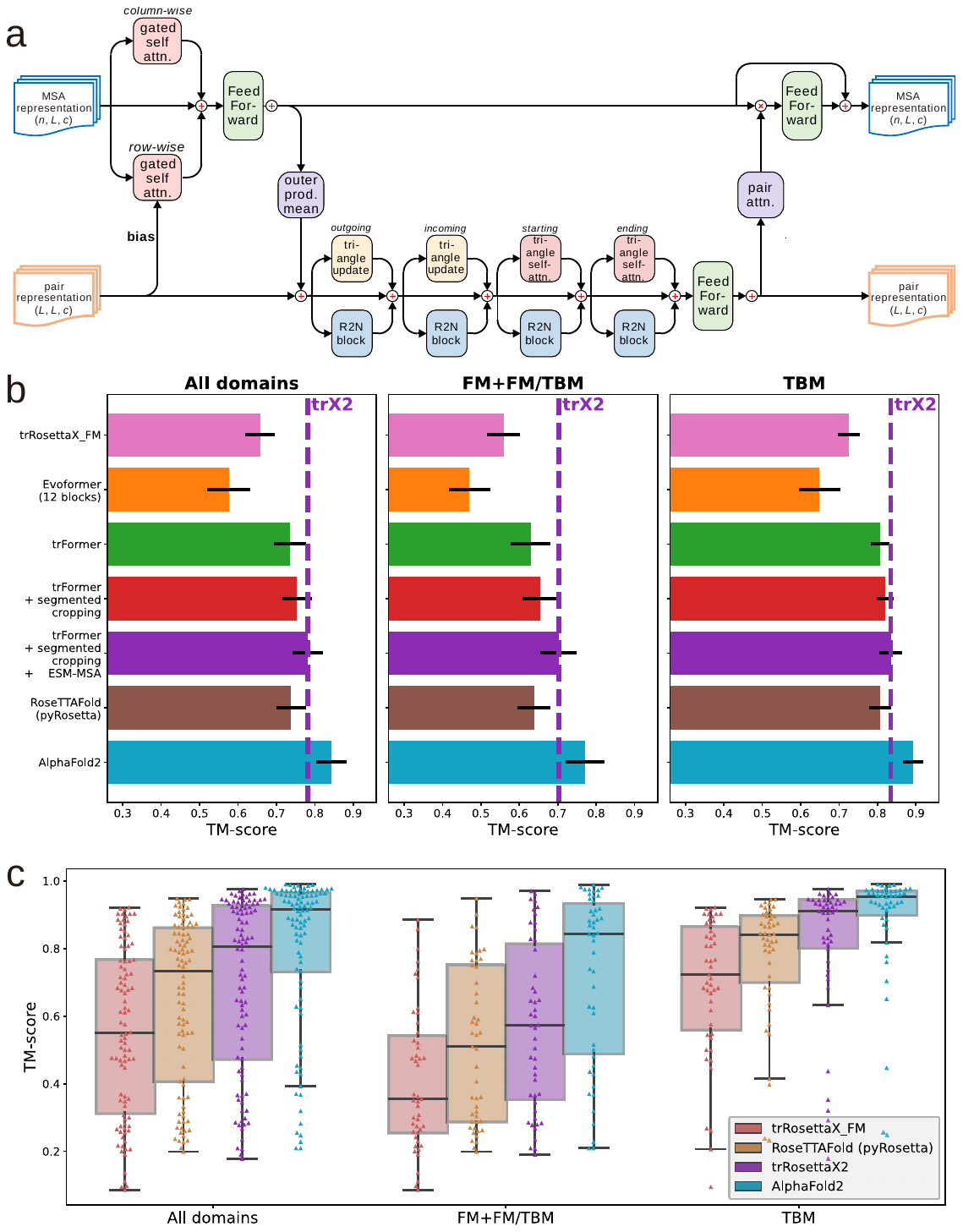

**Fig. S1 | Architecture and performance of trRosettaX2.** **(a)** architecture of a trFormer block. **(b)** average TM-scores on CASP14 domains for trRosettaX_FM, trRosettaX2 and its variants, RoseTTAFold, and AlphaFold2. Error bars represent one-fifth of the standard deviation. **(c)** TM-score distributions on CASP15 domains.

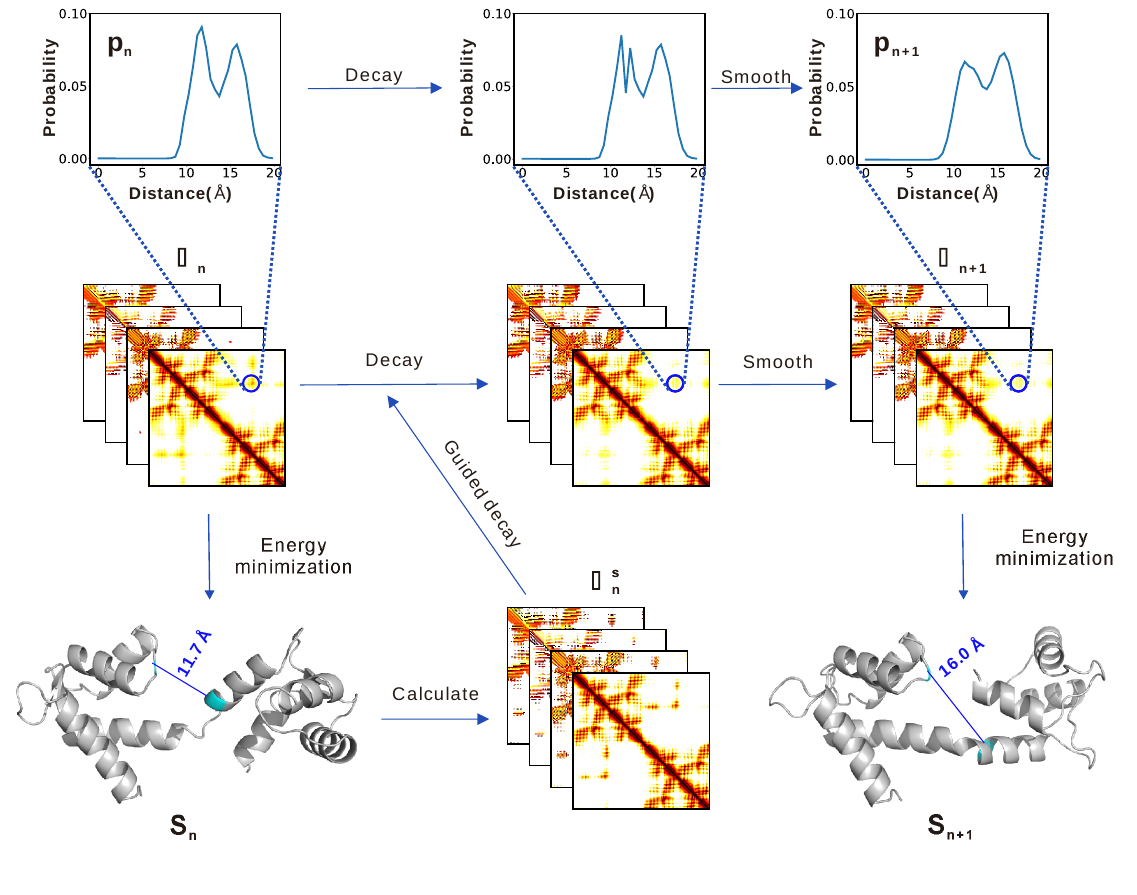

**Fig. S2 | Illustration of one iteration of the heuristic iterative sampling process in trX2-D.**

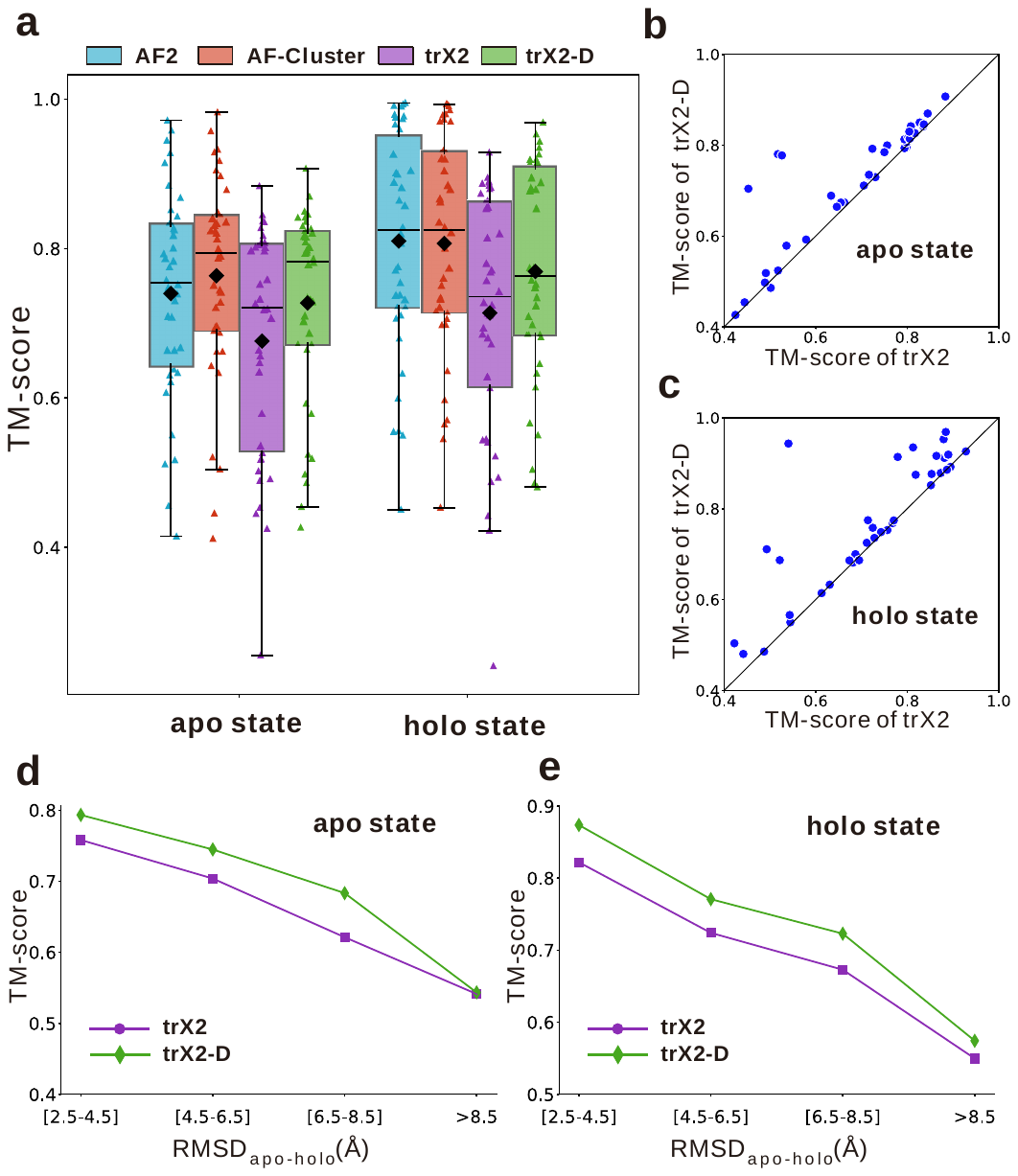

**Fig. S3 | TM-score comparison of trX2-D on proteins with apo and holo states.** **(a)** box plots illustrating the TM-score distribution between predicted and native structures for AF2, AF-Cluster, trX2, and trX2-D, stratified by apo and holo states. Mean TM-scores are indicated by black diamond. **(b, c)** head-to-head TM-score comparison between trX2-D and trX2 for the apo state (b) and holo state (c). **(d, e)** average TM-scores of trX2 and trX2-D predictions for the apo (d) and holo (e) states, grouped by the RMSD_apo-holo_.

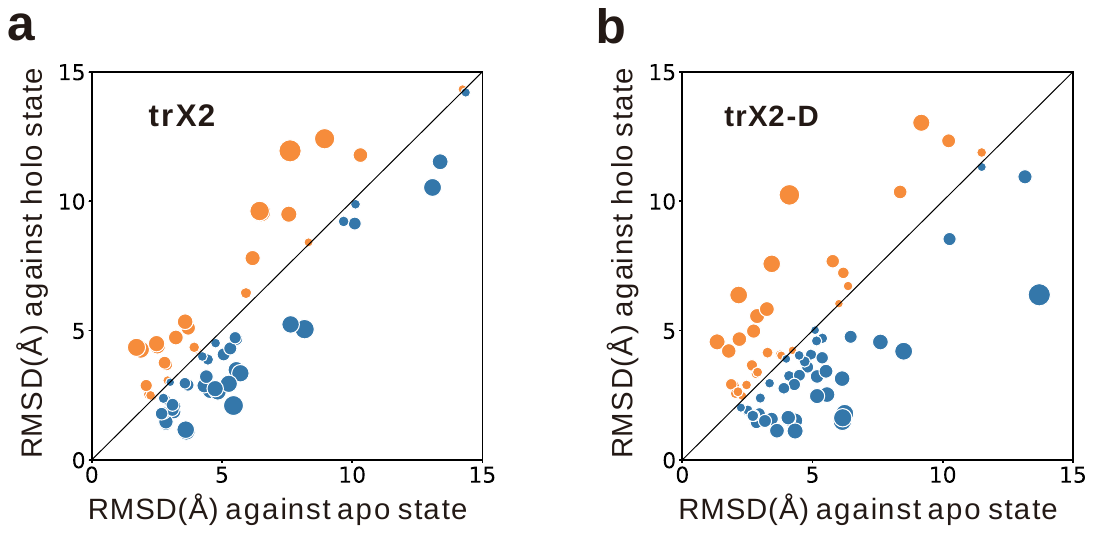

**Fig. S4 | Comparison of RMSDs relative to native apo/holo states for the predicted conformations by trX2 (a) and trX2-D (b).** Point size is proportional to the perpendicular distance to the diagonal.

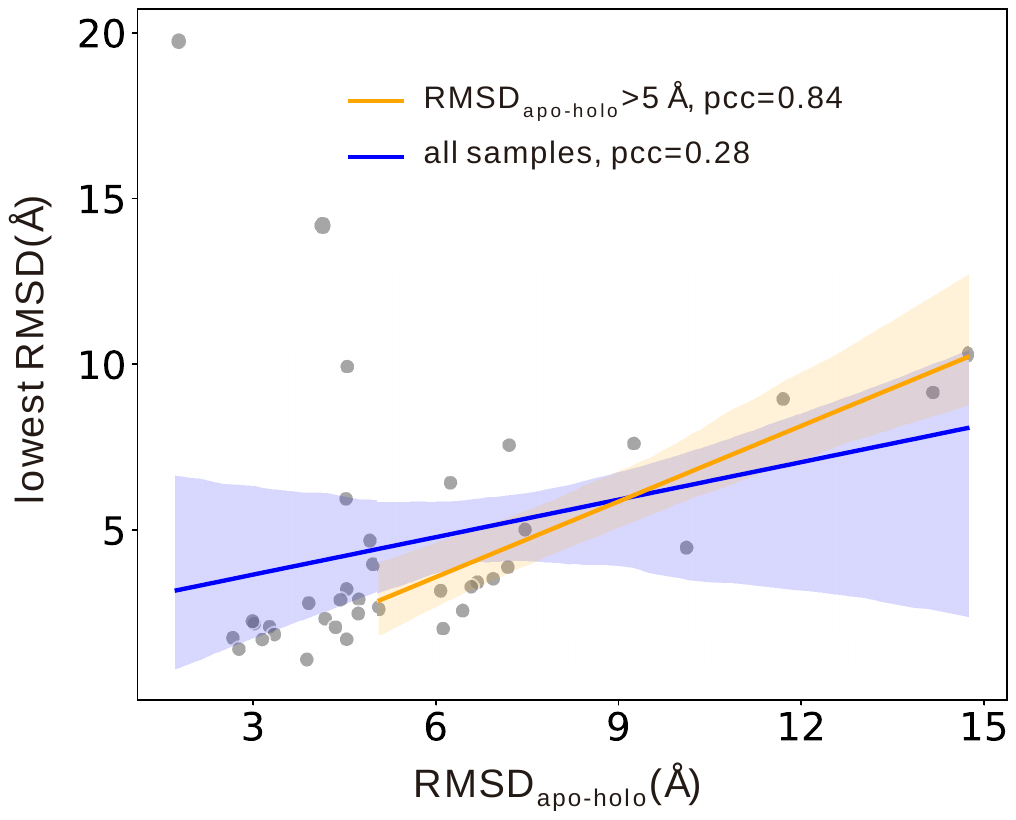

**Fig. S5 | Correlation between native state divergence (RMSD_apo-holo_) and trX2 prediction accuracy (the minimum RMSD to any state).**

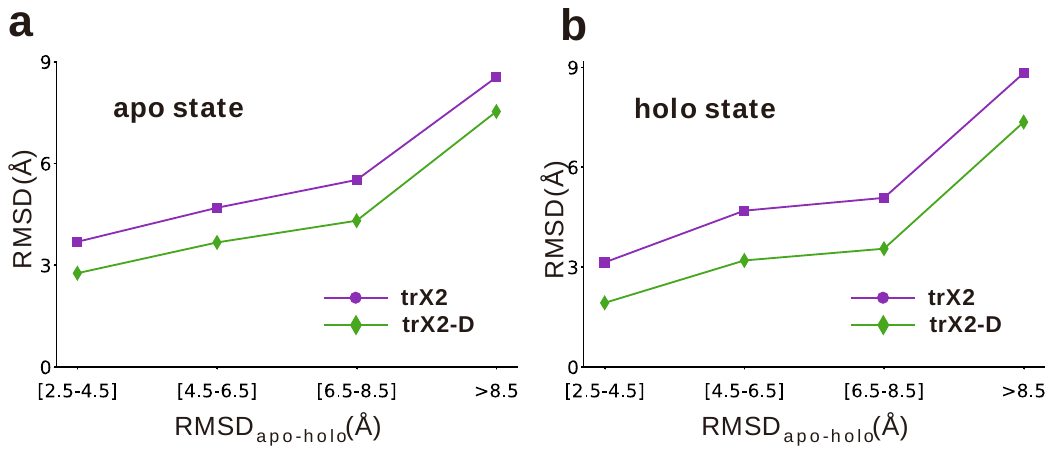
**Fig. S6 | Average RMSDs for trX2 and trX2-D predictions in the apo (d) and the holo (e) states, grouped by the RMSD between the native apo and holo states (RMSD_apo-holo_).** Higher RMSD_apo-holo_ value indicates greater structural divergence between the native apo and holo forms.

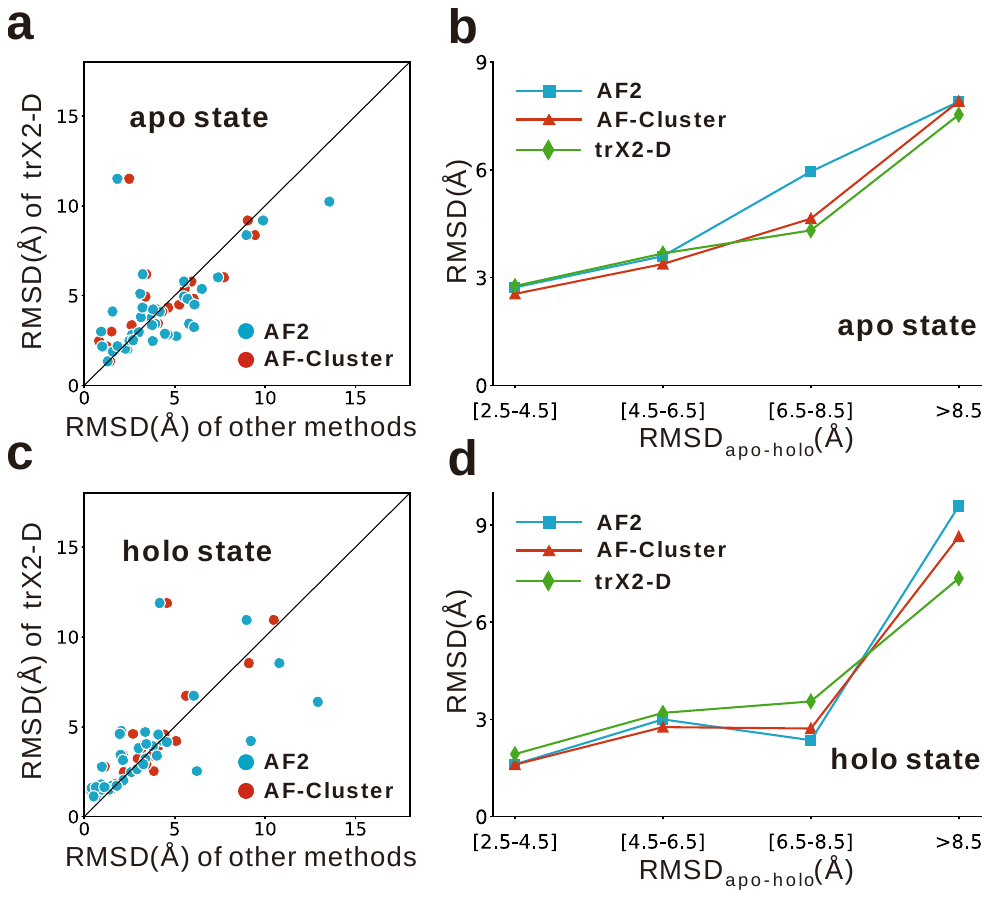

**Fig. S7 | Detailed comparison of prediction accuracy for apo and holo states. (a, c)** head-head RMSD comparison for the apo state (a) and the holo state (c) between trX2-D and other methods (AF2 and AF-Cluster). **(b, d)** the average RMSDs of different methods for the apo state (b) and the holo state (d), grouped by RMSD between native conformations (RMSD_apo-holo_).

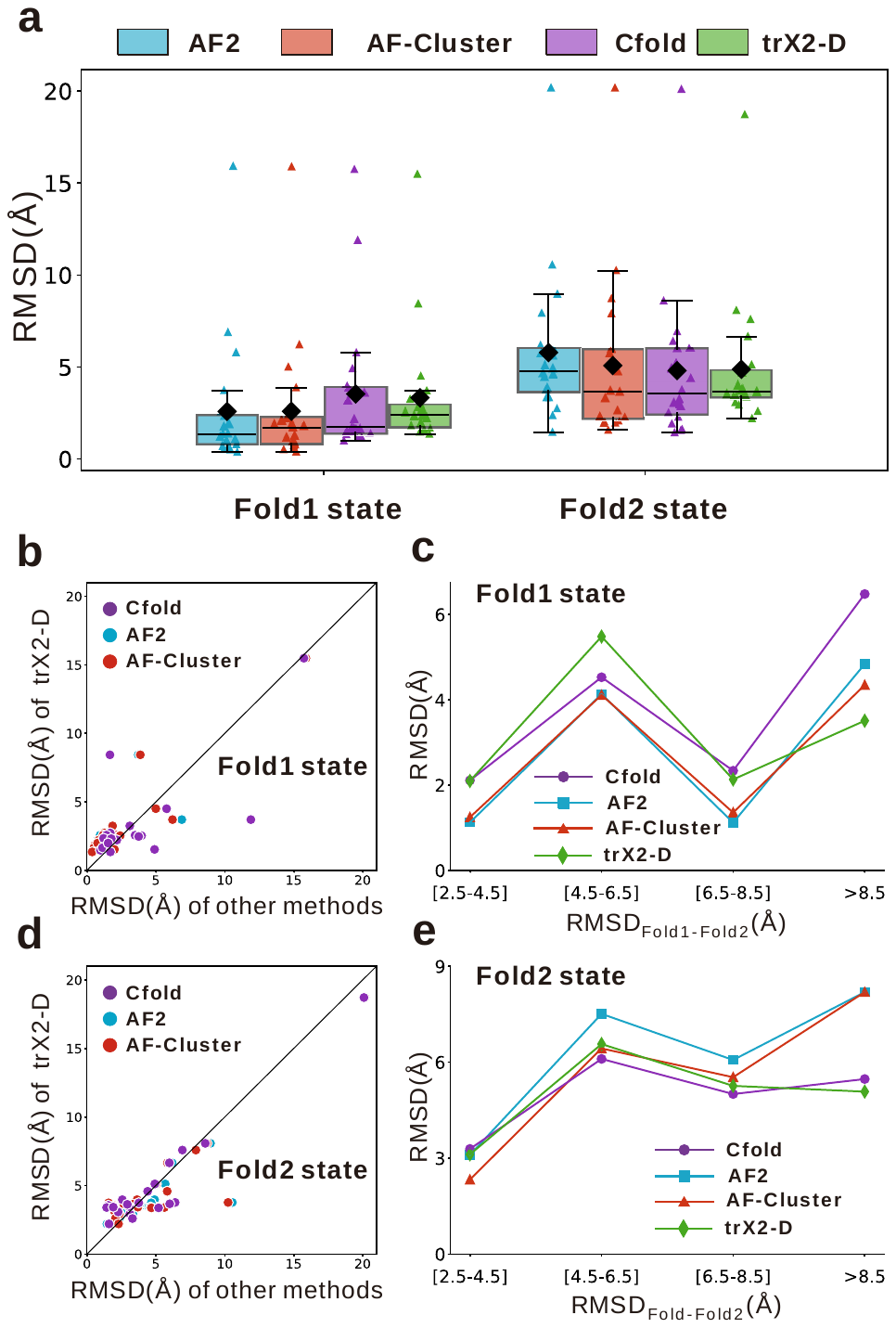

**Fig. S8 | Performance of trX2-D in 20 dual-conformation proteins from the Cfold benchmark.** **(a)** RMSD distributions of predictions from AF2, AF-Cluster, trX2, and trX2-D against native conformations, separated by Fold1 and Fold2 states. Mean RMSDs are indicated by black diamonds. **(b, d)** head-to-head RMSD comparison of trX2-D versus trX2 for the Fold1 (b) and Fold2 (d) states, where points below the diagonal denote trX2-D superiority. **(c, e)** average RMSDs of trX2 and trX2-D predictions for the Fold1 (c) and Fold2 (e) states, grouped by the RMSD between native states (i.e., RMSD_Fold1-Fold2_).

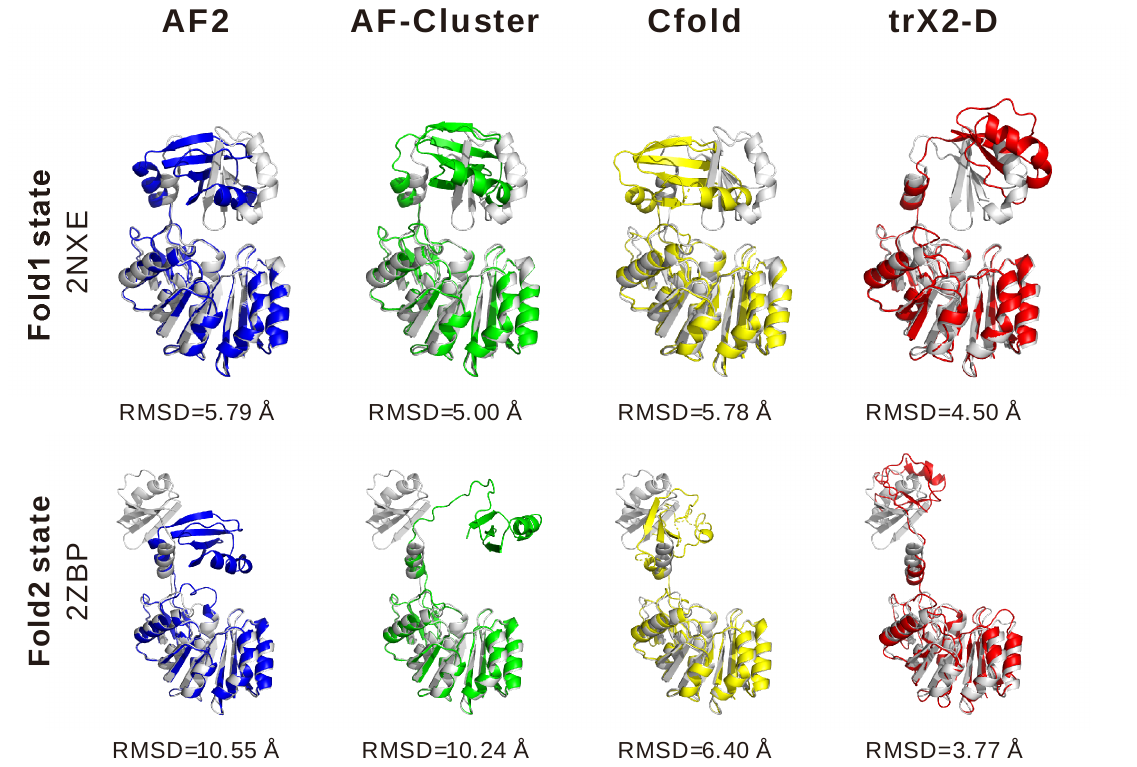

**Fig. S9 | An example in the Cfold benchmark set with a large conformational change.** Fold1 (PDB ID: 2NXE) and Fold2 (PDB ID: 2ZBP) structures, exhibiting notable differences (TM-score_Fold1-Fold2_ = 0.78, RMSD_Fold1-Fold2_ = 11.7 Å), are shown with predictions from AF2, AF-Cluster, Cfold, and trX2-D. trX2-D provides more accurate prediction in both Fold1 and Fold2 states, with conformational difference (TM-score_Fold1-Fold2_ = 0.77, RMSD_Fold1-Fold2_= 11.1 Å) highly close to the native states, and more significant than AF2/AF-Cluster/Cfold (TM-score_Fold1-Fold2_ = 0.95/0.83/0.81, RMSD_Fold1-Fold2_= 1.5/5.4/6.7 Å, respectively).

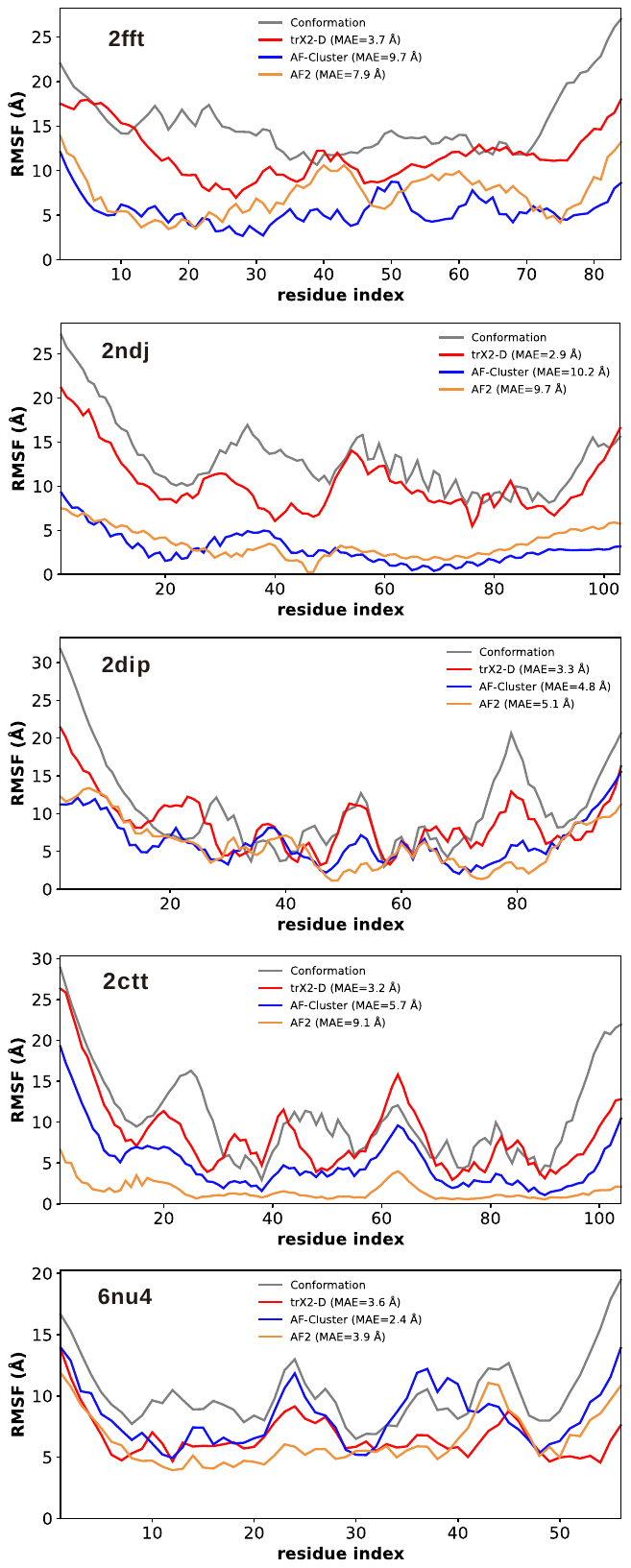

**Fig. S10 (*previous page*) | Per‑residue RMSF profiles for five challenging cases shown in Fig. 6.** Root‑mean‑square fluctuations (RMSF) are plotted against residue index for the native ensemble (grey), AlphaFold2 (orange), AF‑Cluster (blue), and trX2‑Mult (red). Compared to AF2 and AF‑Cluster, trX2‑Mult captures broader structural diversity with RMSF more closely matching the native flexibility.

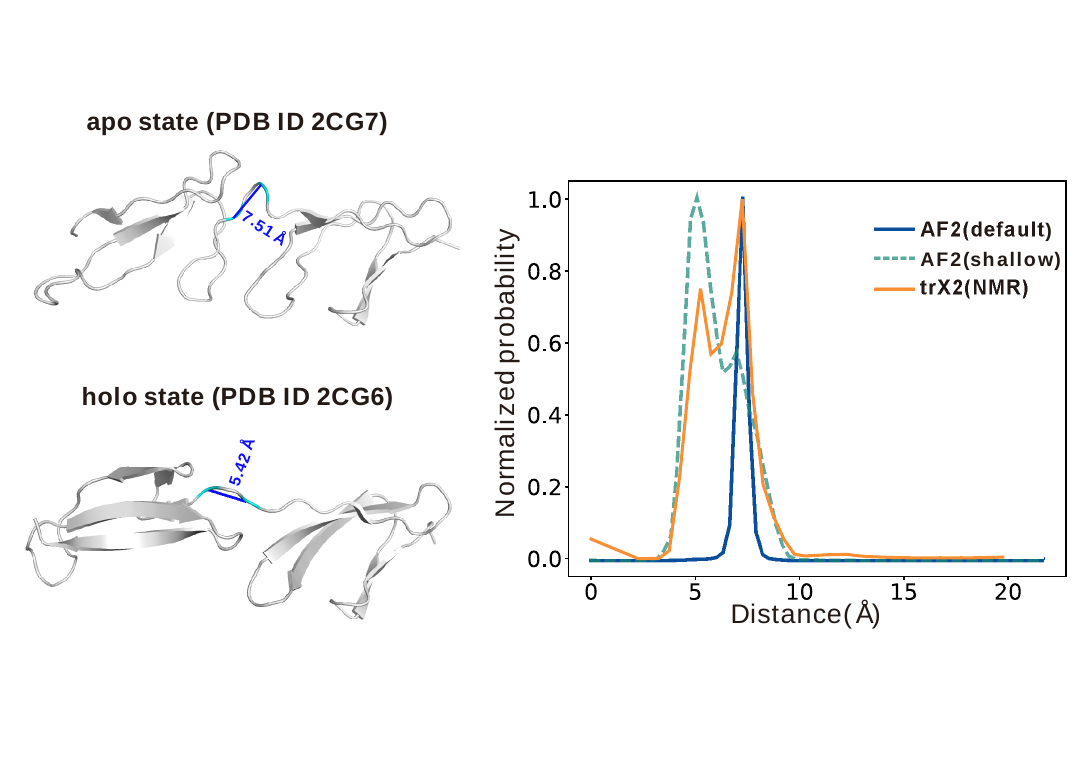
**Fig. S11 | Comparison of distance distributions for a representative residue pair in apo and holo states.** The left panels illustrate the apo (PDB ID 2CG7) and holo (PDB ID 2CG6) structures (cartoon representation), highlighting the residue pair and the corresponding distances in blue. The right panels present the normalized probability distributions of the distance between this pair predicted by trX2(NMR), AF2, and AF2 with subsampled MSA (AF2(shallow)).

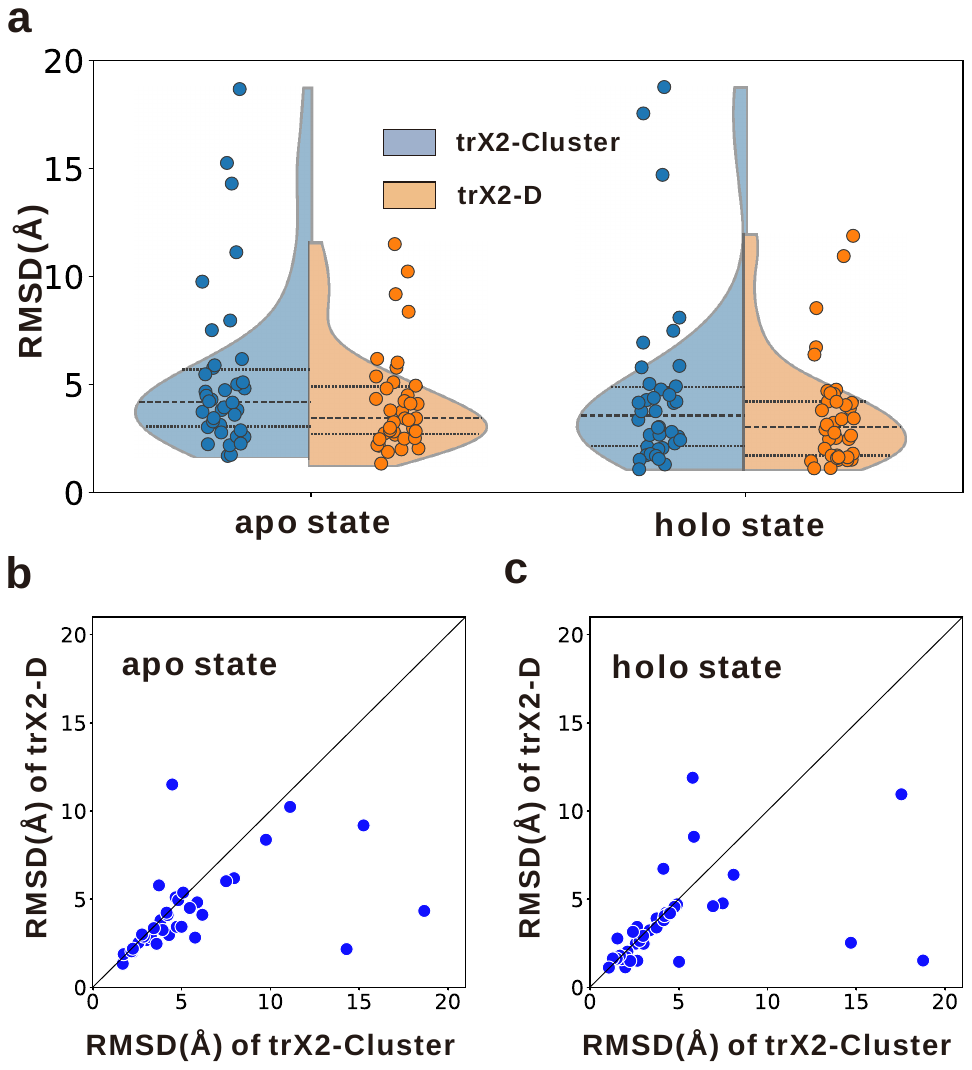

**Fig. S12 | Comparison between trX2-D and trX2-Cluster.** **(a)** violin plots comparing the distribution of RMSD values between predicted and native structures for trX2-Cluster and trX2-D in both apo and holo states. Individual data points are overlaid on the violins. **(b, c)** head-to-head comparisons of RMSD between trX2-D and trX2-Cluster for the apo state (b) and the holo state (c).

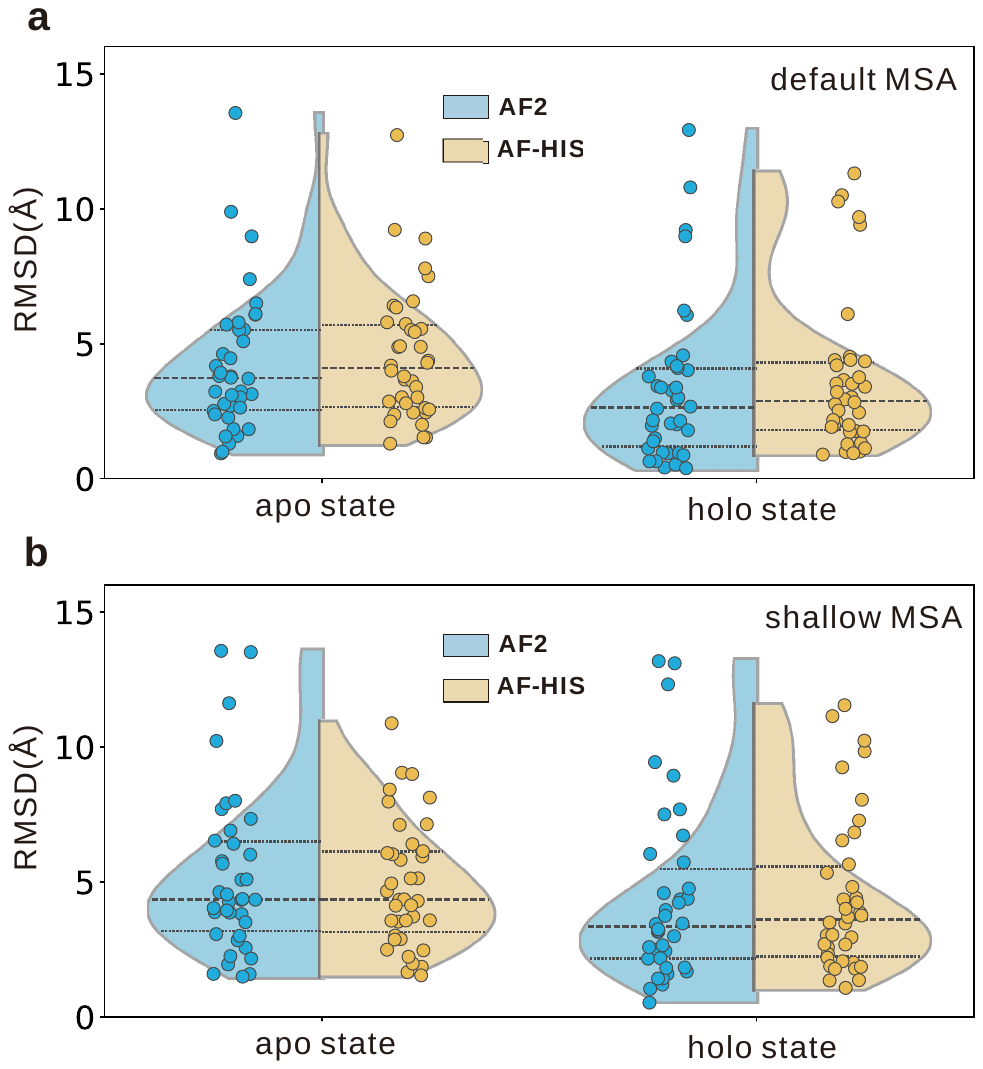

**Fig. S13 | Applying iterative sampling to AF2 with different MSA depths.** **(a)** violin plots showing the distribution of RMSD values for AF2 applying the heuristic iterative sampling strategy (AF-HIS) with default MSA depth, compared to standard AF2 predictions for both apo and holo states. **(b)** Similar violin plots showing the impact of iterative sampling on AF2 predictions generated using shallow MSAs, compared to standard shallow MSA AF2 predictions. Individual data points are overlaid on all plots.

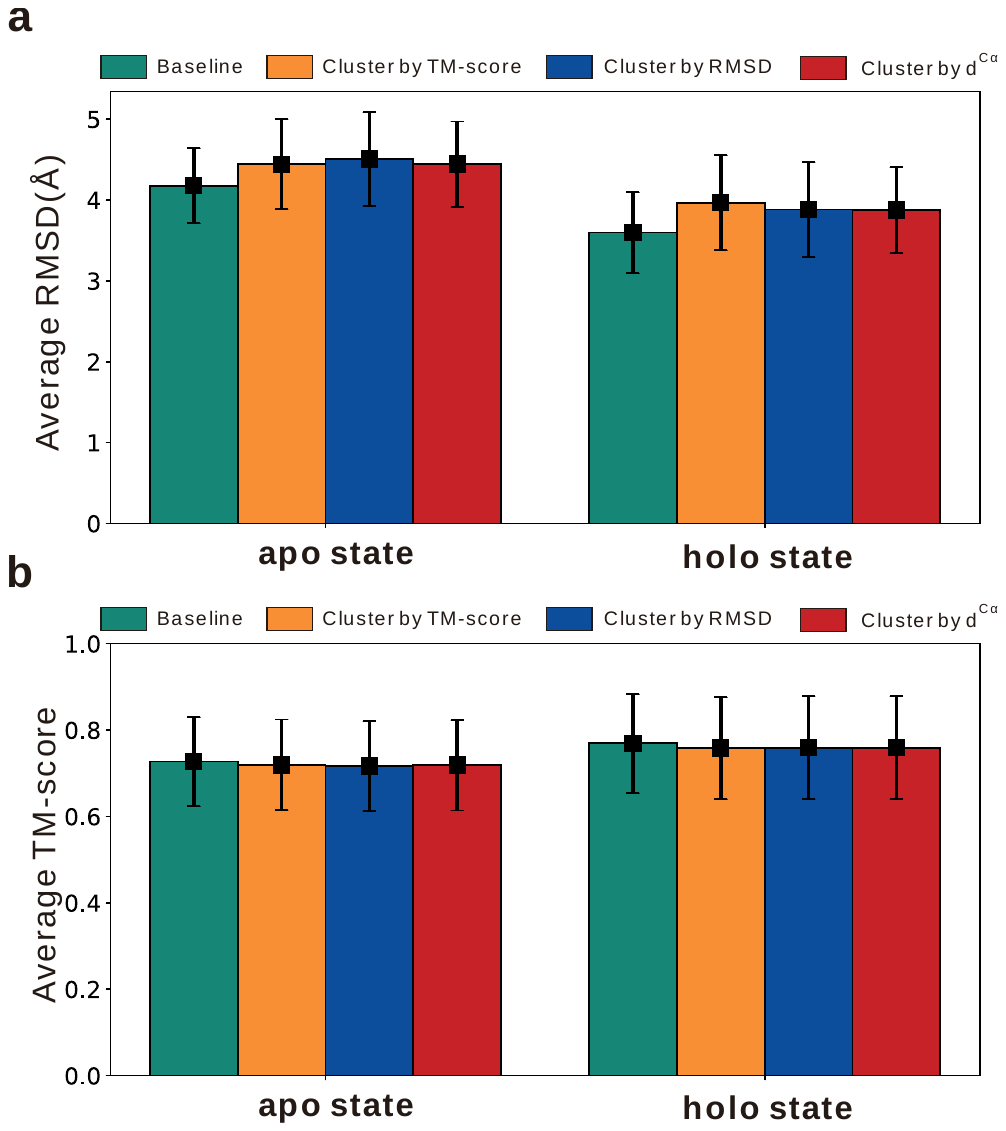

**Fig. S14 | Evaluation of clustering strategies for representative structure selection**. **(a, b)** The average RMSD (a) and TM-score (b) of representative structures selected using k-means clustering based on different structural similarity metrics (TM-score, RMSD, and inter-C$\alpha$ distances $d^{C\alpha}$) for both apo and holo states. A baseline representing the average performance without clustering is also shown. Each error bar represents one-fifth of the standard deviation.

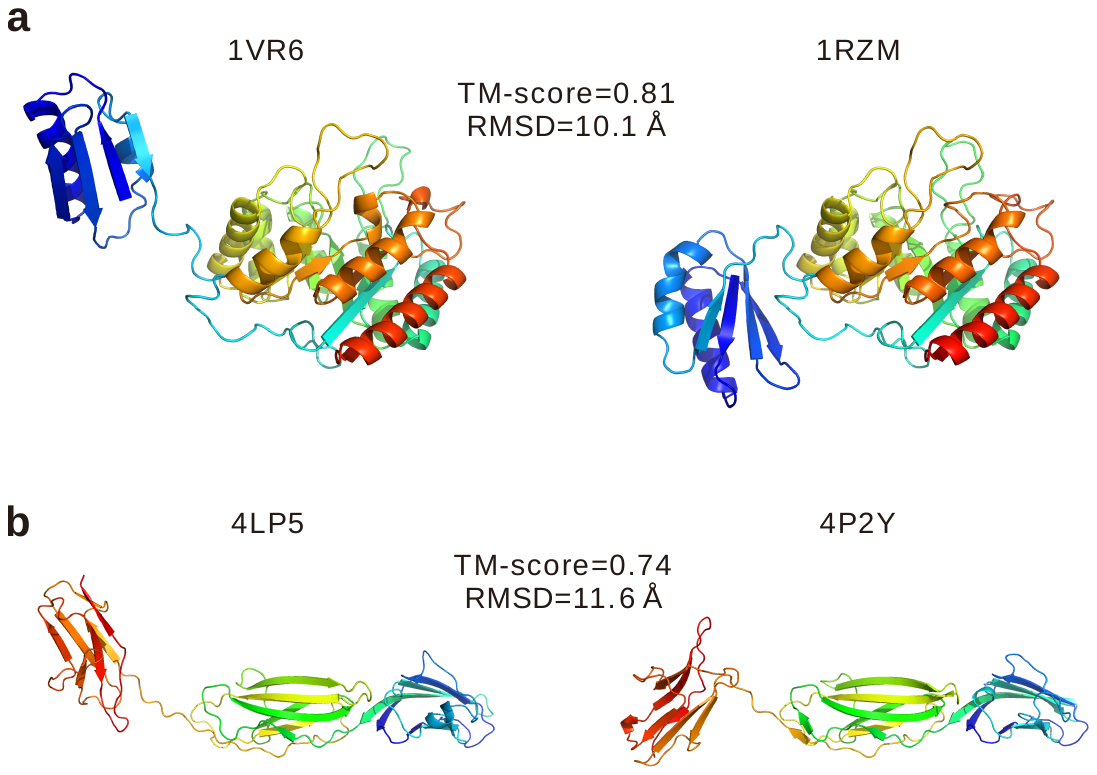

**Fig. S15 | Examples of apo-holo protein pairs exhibiting significant conformational differences despite having high inter-conformation TM-scores.**

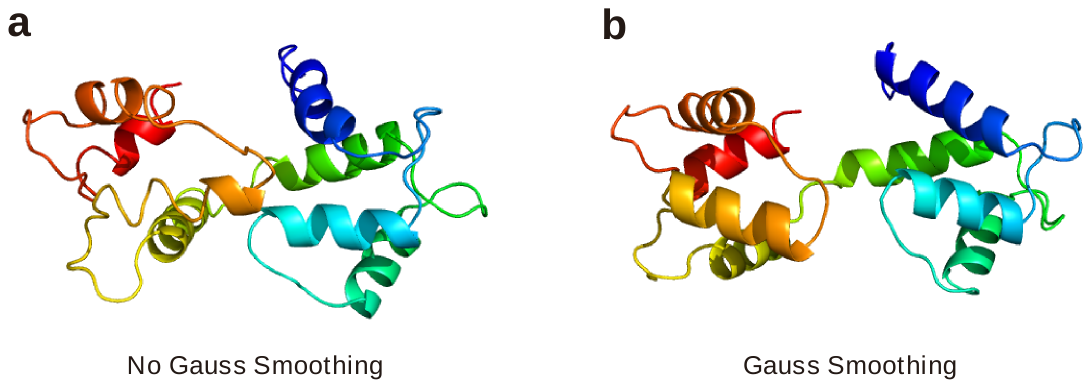

**Fig. S16 | Visualizing the impact of Gaussian smoothing on protein structure representation.** Representative protein structures are shown (a) without and (b), with Gaussian smoothing applied. Gaussian smoothing enhances structural regularity and completeness in the heuristic iterative process.
